# Use of a pandemic H1N1 strain with updated haemagglutinin and neuraminidase results in increased nasopharyngeal shedding and improved immunogenicity to Russian-backbone live attenuated influenza vaccine among children aged 2 – 4 years old: an open-label, prospective, observational, phase 4 study in The Gambia

**DOI:** 10.1101/519256

**Authors:** Benjamin B. Lindsey, Ya Jankey Jagne, Edwin P. Armitage, Anika Singanayagam, Hadijatou J. Sallah, Sainabou Drammeh, Elina Senghore, Nuredin I. Mohammed, David Jeffries, Katja Höschler, John S. Tregoning, Adam Meijer, Ed Clarke, Tao Dong, Wendy Barclay, Beate Kampmann, Thushan I. de Silva

**Author notes:** These authors contributed equally to the work (Benjamin B. Lindsey, Ya Jankey Jagne, Edwin P. Armitage). Corresponding author: Thushan I. de Silva.

## Abstract

**Background:** Poor efficacy and effectiveness of the pandemic H1N1 (pH1N1) component in intranasal live attenuated influenza vaccine (LAIV) has been demonstrated in several studies. The reasons for this are unclear, but may be due to impaired replicative fitness of pH1N1 A/California/07/2009-like (Cal09) LAIV strains.

**Methods:** In an open-label, prospective, observational, phase 4 study, we evaluated the impact of updating the pH1N1 component in the Russian-backbone trivalent LAIV from Cal09 in 2016-17 (n=118) to an A/Michigan/45/2015-like strain (A/17/New York/15/5364, NY15) in 2017-18 (n=126), on shedding and immunogenicity in Gambian children aged 2-4 years old. The study was nested within a larger randomised controlled trial investigating LAIV-microbiome interactions (ClinicalTrials.gov NCT02972957).

**Findings:** Cal09 showed impaired nasopharyngeal shedding compared to H3N2 and influenza B, along with sub-optimal serum antibody and T-cell responses. Following the switch to NY15, a significant increase in pH1N1 shedding was seen, along with improvements in seroconversion and influenza-specific CD4+ T-cell responses. Viral kinetics *in vitro* mirrored these findings, with NY15 showing greater replicative ability than Cal09 in human nasal epithelial cells. Persistent shedding to day 7 was independently associated with both seroconversion and CD4+ T cell response in multivariable logistic regression.

**Interpretation:** Our results suggest that the pH1N1 component switch in LAIV may have overcome problems in prior formulations. LAIV effectiveness against pH1N1 should therefore improve in upcoming influenza seasons. Our data also highlight the importance of evaluating replicative fitness, in addition to antigenicity, when selecting annual LAIV components and design of potentially more effective vaccines.

**Funding:** The Wellcome Trust.

## Introduction

There are concerns regarding the protection against pandemic H1N1 (pH1N1) influenza from live attenuated influenza vaccine (LAIV). Reduced effectiveness against pH1N1 of LAIV compared to inactivated influenza vaccine (IIV) in the United States (US) (Ann Arbor-backbone LAIV; Medimmune) since 2013-14 resulted in the Advisory Committee on Immunisation Practices removing their recommendation for LAIV use in 2016.^1^ A randomised controlled trial (RCT) of Russian-backbone LAIV (Nasovac-S, Serum Institute of India Pvt Ltd) among children aged 2 – 5 years in Senegal also failed to demonstrate efficacy (0.0%, 95% confidence interval (CI) −26.4-20.9)in 2013, when pH1N1 was the predominant circulating vaccine-matched virus.^2^ In these studies, both formulations contained haemagglutinin (HA) and neuraminidase (NA) from pH1N1 A/California/07/2009-like (Cal09) viruses. Why protection conferred by the pH1N1 component in LAIV has been suboptimal is unclear. Potential reasons include pre-existing immunity, poor viral replicative fitness or competition from other co-formulated strains, all limiting pH1N1 ‘take’ and immunogenicity.^1^

The findings in Senegal are particularly pertinent as the burden of influenza in Africa is high, with influenza-related hospitalisations in children <5 years approximately three-fold higher than in Europe.^3^ Previously demonstrated superior efficacy of LAIV over IIV in young children (predominantly in high-income settings), needle-free delivery and lower manufacturing costs make LAIV an attractive option to tackle this burden in Africa.^1^ There are, however, limited LAIV studies in African cohorts and no published immunogenicity data from African children to date.^4^ In particular, the absence of immunological endpoints from the RCT in Senegal make it difficult to understand the reasons for the lack of efficacy observed.^2^

To better understand how differences in strain shedding and immunogenicity may explain these findings, we immunised one cohort of influenza vaccine-naïve Gambian children with the Russian-backbone LAIV 2016-17 formulation, followed by a second cohort with the 2017-18 formulation of the same vaccine. In 2017-18, the pH1N1 Cal09 strain (A/17/California/2009/38) was updated according to WHO recommendations to an A/Michigan/45/2015-like strain (A/17/New York/15/5364, NY15), following antigenic drift. This first ever recommended update to pH1N1 provided us with a unique opportunity to compare replicative ability and immunogenicity of these two pH1N1 strains.

## Methods

### Study design and participants

An open-label, prospective, observational, phase 4 immunogenicity study was conducted in Sukuta, a periurban area in The Gambia. This study was nested within a larger RCT (ClinicalTrials.gov NCT02972957) comparing microbiome changes in LAIV and unvaccinated arms, and represents data from all enrolled children given LAIV to measure shedding and immunogenicity endpoints. Following community sensitisation, parents expressing an interest in the study were invited for consent discussions. Eligible children had to be aged 24-59 months and clinically well, with no history of respiratory illness within the last 14 days (full criteria in appendix).

The study was approved by the Gambia Government/MRC Joint Ethics Committee and the Medicines Control Agency of The Gambia, and conducted according to ICH-GCP standards. A parent provided written or thumb-printed informed consent for their children to participate. Where parents were not English literate, an impartial witness was present throughout the informed consent discussion undertaken in a local language and signed to confirm completeness of the consent provided.

### Procedures

Children received one dose of intranasal trivalent Russian-backbone LAIV (Nasovac-S, Serum Institute of India Pvt Ltd) northern hemisphere (NH) formulation in either 2017 (2016-17 NH formulation) or 2018 (2017-18 NH formulation). In the 2017-18 LAIV, HA and NA from pH1N1 Cal09 were replaced with those from NY15, while identical H3N2 and B/Victoria lineage (B/Vic) strains were used. Vaccine titres per dose (50% Egg Infectious Doses (EID50)/ml) were 1×10^8.0^ for pH1N1, 1×10^7.5^ for H3N2, 1×10^7.2^ for B/Vic in the 2016-17 LAIV and 1×10^7.7^ for pH1N1, 1×10^7.6^ for H3N2, 1×10^7.3^ for B/Vic in the 2017-18 LAIV.

Nasopharyngeal swabs (NPS) were taken pre-vaccination (day 0, D0), at day 2 (D2) and day 7 (D7) using flocked swabs (Copan FloQSwabs™). Oracol+ swabs (Malvern Medical Development Ltd.) were used to collect buccal cavity oral fluid (OF) at D0 and day 21 (D21). Whole blood was collected for flow cytometry and serum separation at D0 and D21. D21 was chosen to measure vaccine response in line with previously published studies.^5–7^ NPS, OF and serum samples were stored at −70°C prior to further processing.

Haemagglutinin inhibition (HAI) assays were performed according to standard methods, using vaccine HA- and NA-matched viruses.^8^ Seroconversion was defined as a ≥4-fold titre increase (to ≥1:40) from D0 to D21. Total and influenza HA-specific immunoglobulin A (IgA) in OF was detected using a previously described enzyme-linked immunosorbent assay, using recombinant vaccine-matched HA.^9^ Samples were assayed at dilutions of 1:1000 – 1:20000 for total IgA and neat to 1:16 for influenza-specific IgA, and quantified using an IgA standard curve. Neat samples with influenza-specific IgA below the limit of quantitation (LOQ) were assigned LOQ values. D0 to D21 fold change in proportion of influenza-specific:total IgA was calculated. A 2-fold increase was considered a significant response.^10^

T-cell responses were quantified by stimulating fresh whole blood (200μl) for 18 hours with overlapping 15-18-mer peptide pools (2μg/ml) covering vaccine-matched whole HA, Matrix and Nucleoprotein (MNP), along with co-stimulatory antibodies anti-CD28 and anti-CD49 (BD Biosciences). Influenza B responses were measured in 2018 only. Intracellular cytokine staining for IFN-γ and IL-2 was performed and cells analyzed with an LSR Fortessa flow cytometer (table S2, figures S2 and S3).^11^ Responses in the negative (anti-CD28/anti-CD49) controls were subtracted from peptide-stimulated conditions prior to further analysis. Negative values were set to zero. To avoid systematic bias in doing so, a threshold (table S3, based on the distribution of negative values) was set below which all positive values were also considered a non-response, as described previously.^11,12^ In analyses calculating D0 to D21 fold-change, null responses were assigned a value half-way between zero and this threshold. A 2-fold increase post-LAIV was considered a significant response.

Vaccine shedding was assessed with monoplex reverse-transcriptase polymerase chain-reaction (RT-PCR) using HA-specific primers and probes (appendix, table S4). In 2018, fully quantitative RT-PCR (qRT-PCR) results were obtained by inclusion of a standard curve with known log10 50% Egg Infectious Doses/ml (EID/ml)(appendix, figure S4). RT-PCR assays with primers and probes mapping to internal genes were used to distinguish LAIV from seasonal influenza viruses (appendix, table S4).^13^ Despite optimisation of assay conditions, maximum LAIV dilutions detected by LAIV-specific RT-PCR were at least one log10 lower than those detected by HA-specific RT-PCR (Table S5). Only samples with ct values of ≤30 in seasonal influenza assays were therefore tested with LAIV-specific assays, with 100% confirmed as LAIV strains.

Primary human nasal epithelial cell (hNEC) cultures (Mucil Air™, Epithelix Sàrl) were used for *in vitro* viral replication experiments. Madin-Darby Canine Kidney (MDCK) cells (ATCC) and MDCK-SIAT cells (WHO CC London) were maintained at 37°C with 5% CO2 in Dulbecco’s modified Eagle’s Medium (DMEM) (Gibco-Life technologies) supplemented with 10% foetal bovine serum, 1% penicillin/streptomycin and 1% non-essential amino acids. 1mg/mL G418 (Gibco-Life technologies) was also added for MDCK-SIAT cells. Viral stocks of Nasovac-S monovalent forms were titrated by plaque assay performed at 32°C on MDCK cells (for pH1N1 and influenza B) or MDCK-SIAT (for H3N2) cells. Apical surfaces of hNECs were inoculated with each monovalent virus (multiplicity of infection 0.01 plaque forming units (PFU)/cell) for 1 hour at 32°C, 5% CO2 in triplicate. The inoculum was removed and the apical surface washed with DMEM before incubating at 32°C. At indicated time points, DMEM was added, incubated for 30 minutes, removed and stored, then later titrated by plaque assay. Experiments were performed on two separate occasions using cells from different donors.

To assess acid-stability of pH1N1 strains, Cal09 or NY15 were mixed with pH-adjusted MES buffers (100mM MES, 150mM NaCl, 0.9mM CaCl_2_, 0.5mM MgCl_2_) in triplicate (1:10 dilution) and incubated for 15 minutes at room temperature. The buffer was inactivated with DMEM and infectious virus titrated by plaque assay.

### Outcomes

The primary shedding and immunogenicity outcomes were the percentage of children with LAIV strain shedding at D2 and D7, HAI seroconversion and increase in influenza HA-specific IgA and T-cell responses at D21 post-LAIV.

### Statistical analysis

Differences in proportions between years were examined using Chi-squared or Fisher’s exact test and continuous variables between years using the unpaired t-test or Mann-Whitney U test. Differences in viral load between strains were examined using the Friedman test (with Dunn’s post-test for multiple comparisons). Pairwise viral load correlations were assessed using Spearman’s rank-order correlation. Wilcoxon signed-rank test was used to compare T-cell responses before and after vaccination. Relationships between shedding and other variables was assessed using logistic regression. Viral loads were determined from standard curves using Python 3.6 (SciPy package).^14^ The proportion of mono- and dual-functional T-cell responses were estimated using Boolean gating on FlowJo 10.4^15^ and statistical significance between timepoints tested with the Permutation test in SPICE V6.0.^12^ Proportions are displayed with 95% confidence intervals. Sample size recruited was based on LAIV-microbiome endpoints not presented in this paper (appendix). All tests were two-sided at 5% significance level and were Bonferroni-adjusted for multiple comparisons within each set of analyses. Statistical analyses were carried out using R^16^, Stata12^17^ and GraphPad Prism 5.0d.^18^

## Results

Between February and April 2017, 118 children were enrolled and received one dose of 2016-17 NH formulation LAIV (figure 1A). Between January and April 2018, a separate cohort of 135 children were enrolled and received one dose of 2017-18 NH formulation LAIV (figure 1B). The study was conducted outside the peak influenza transmission season based on surveillance data from Senegal.^19^ 118 children in 2017 and 126 children in 2018 completed the study. No significant difference in baseline demographics was seen between the two cohorts, other than baseline HAI titres (Table 1).

**Figure 1.**
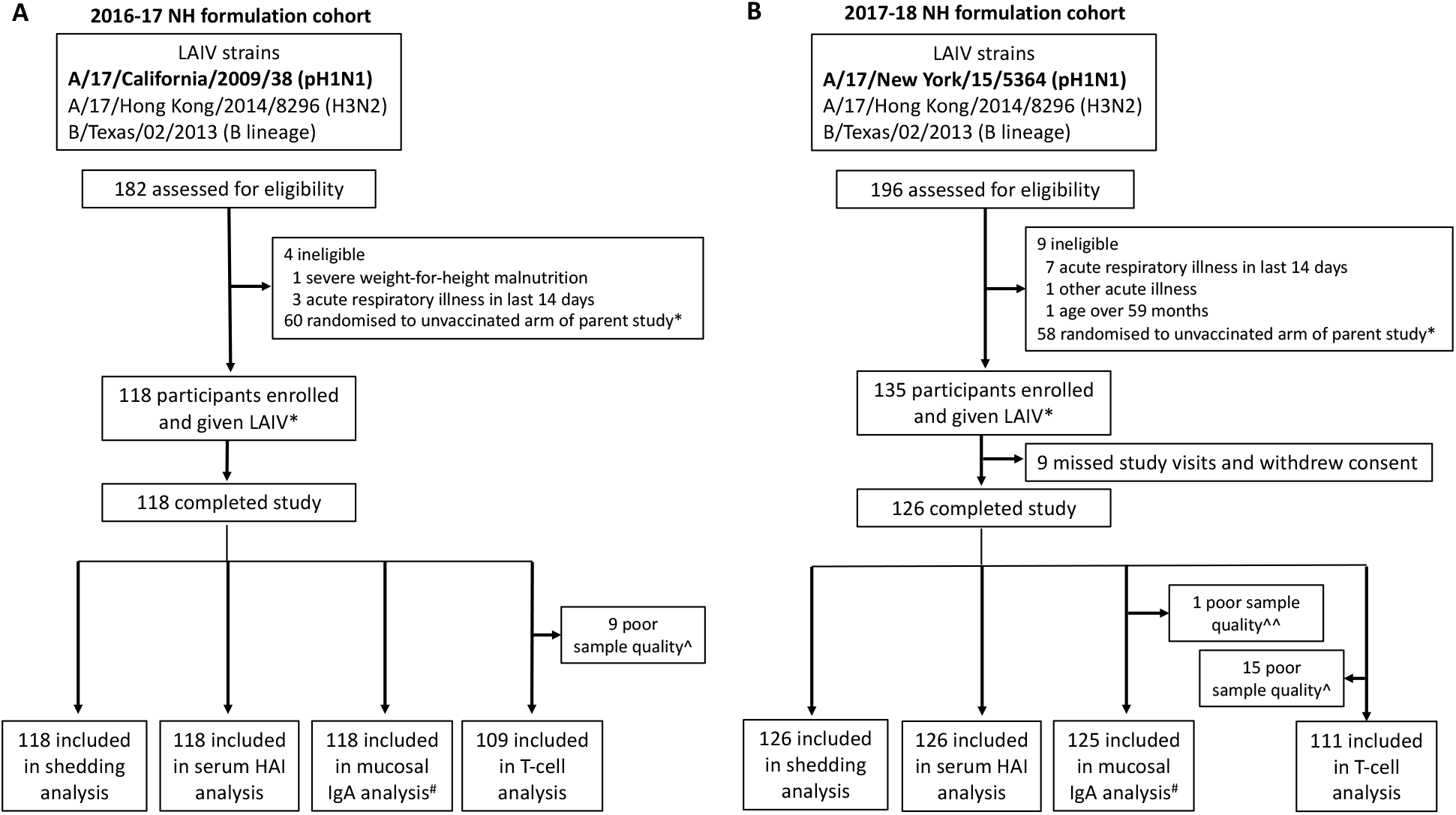
Study profile. Overview of participants who received the 2016-17 northern hemisphere (NH) Russian-backbone LAIV formulation **(A)** and 2017-18 NH formulation **(B)**. All visits were within protocol-defined windows: +1 day for day 2 (D2) visit, +7 days for day 7 (D7) visits and +7 days for day 21 (D21) visits. In the 2016-17 cohort, all (118/118) D2 visits were 2 days post-LAIV, 115/118 of D7 visits were 7 days post-LAIV (3 were 8, 12 and 14 days post-LAIV) and 112/118 of D21 visits were 21 days post-LAIV (5 were 22 days post-LAIV and 1 was 25 days post-LAIV). In the 2017-18 cohort, 122/126 D2 visits were 2 days post-LAIV (4 were 3 days post-LAIV), 119/126 D7 visits were 7 days post-LAIV (7 were 8 days post-LAIV) and 117/126 of D21 visits were 21 days post-LAIV (8 were 22 days post-LAIV and 1 was 26 days post-LAIV). *The study was nested within a larger randomized controlled trial (ClinicalTrials. Gov identifier NCT02972957, see Supplementary Information, Figure S1 and Table S1). ^sparse cell populations seen when acquisition on flow cytometer. ^^Total Immunoglobulin A (IgA) not detected in sample. ^#^No pH1N1 data for 1 sample in 2016-17 cohort, no pH1N1 data for 4 samples and H3N2 data for 3 samples in 2017-18 cohort due to inadequate sample volume. HAI = haemagglutination inhibition.

**Table 1.**
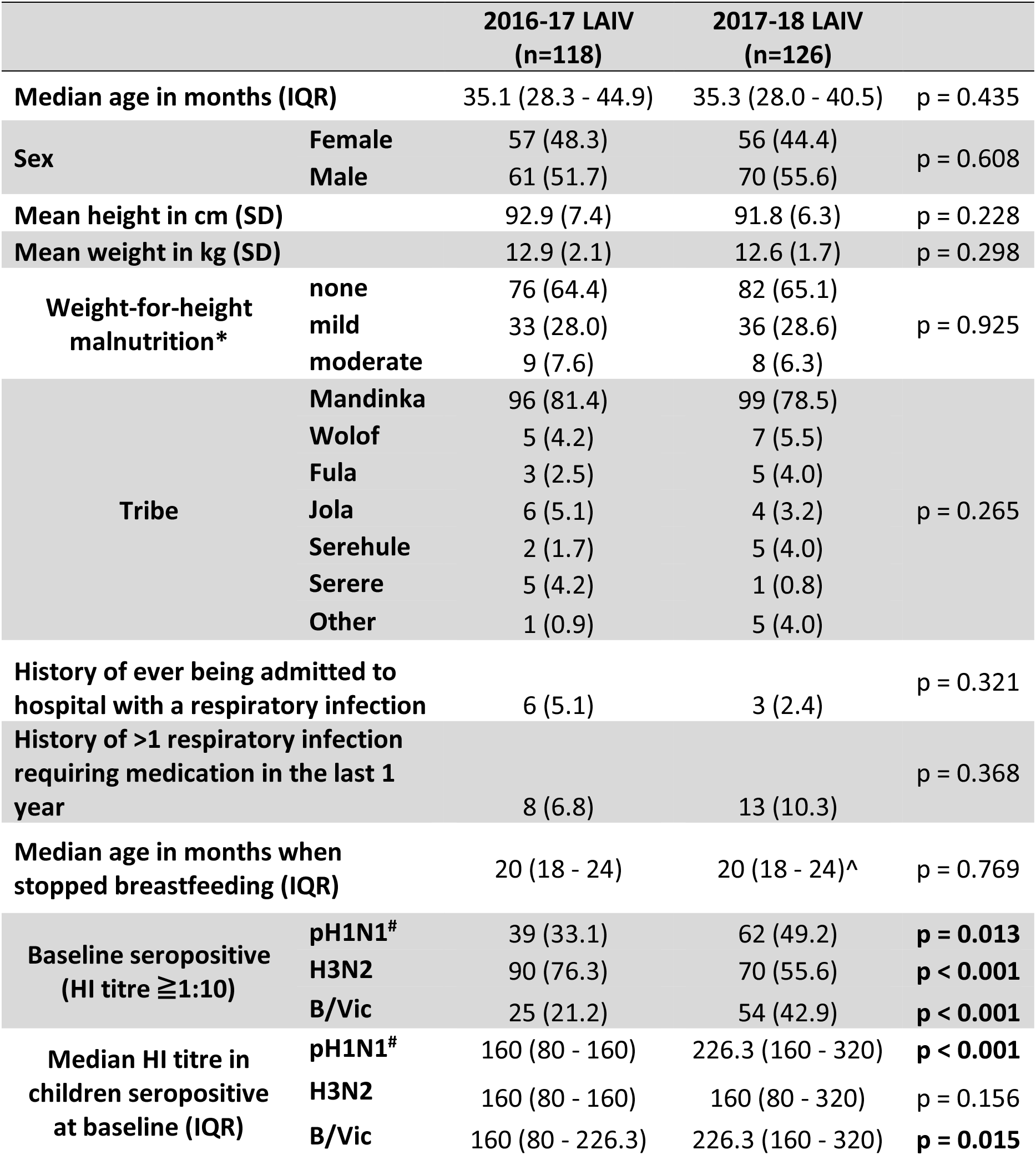
Demographic characteristics and baseline influenza serology data from participants included in the two LAIV cohorts. Data are n (%) unless otherwise stated. IQR = inter-quartile range, SD = standard deviation, HI = haemagglutination inhibition. Malnutrition was categorised based on weight-for-height SD (Z score): none (> −1), mild (−2 to < −1), moderate (−3 to < −2). *children with severe malnutrition (weight-for-height SD < −3) were excluded. ^missing data from 2 children. ^#^Pandemic H1N1 virus used for serum haemagglutination inhibition assays was changed for the cohort given 2017-18 LAIV to reflect the update from Cal09 to NY15.

No influenza strains were detected from NPS taken immediately prior to vaccination in any children. After the 2016-17 LAIV, pH1N1 shedding was seen in significantly fewer children (13.6%, 8.0-21.1) at D2 compared to H3N2 (45.8%, 36.6-55.2, p<0.001) or B/Vic (80.5%, 72.2-87.2, p<0.001). No pH1N1 shedding was seen at D7, with H3N2 shedding seen in 17.8% (11.4-25.9)and B/Vic shedding seen in 59.3% (49.9-68.3). The 2017-18 LAIV resulted in significantly more children shedding pH1N1 at D2 (63.5%, 54.4-71.9, p<0.001), with shedding now detected in 51.6% (42.5-60.6) at D7 (figure 2A). Significantly higher pH1N1 nasopharyngeal viral loads (lower ct values) were also seen with the 2017-18 compared to the 2016-17 LAIV at D2 (figure 2B). Quantitative RT-PCR data showed that NY15 viral loads were in fact now significantly higher than H3N2 and B/Vic at D2 (figure 2C). We explored whether the improved replication of pH1N1 with the 2017-18 LAIV resulted in greater competition with H3N2 and B/Vic (and therefore lower viral loads of these strains), by comparing viral loads in co-shedders of each. No negative impact on H3N2 and B/Vic replication was seen, with a significant positive correlation between pH1N1 and H3N2 shedding seen at D2 and D7 (figures 2D – 2G).

**Figure 2.**
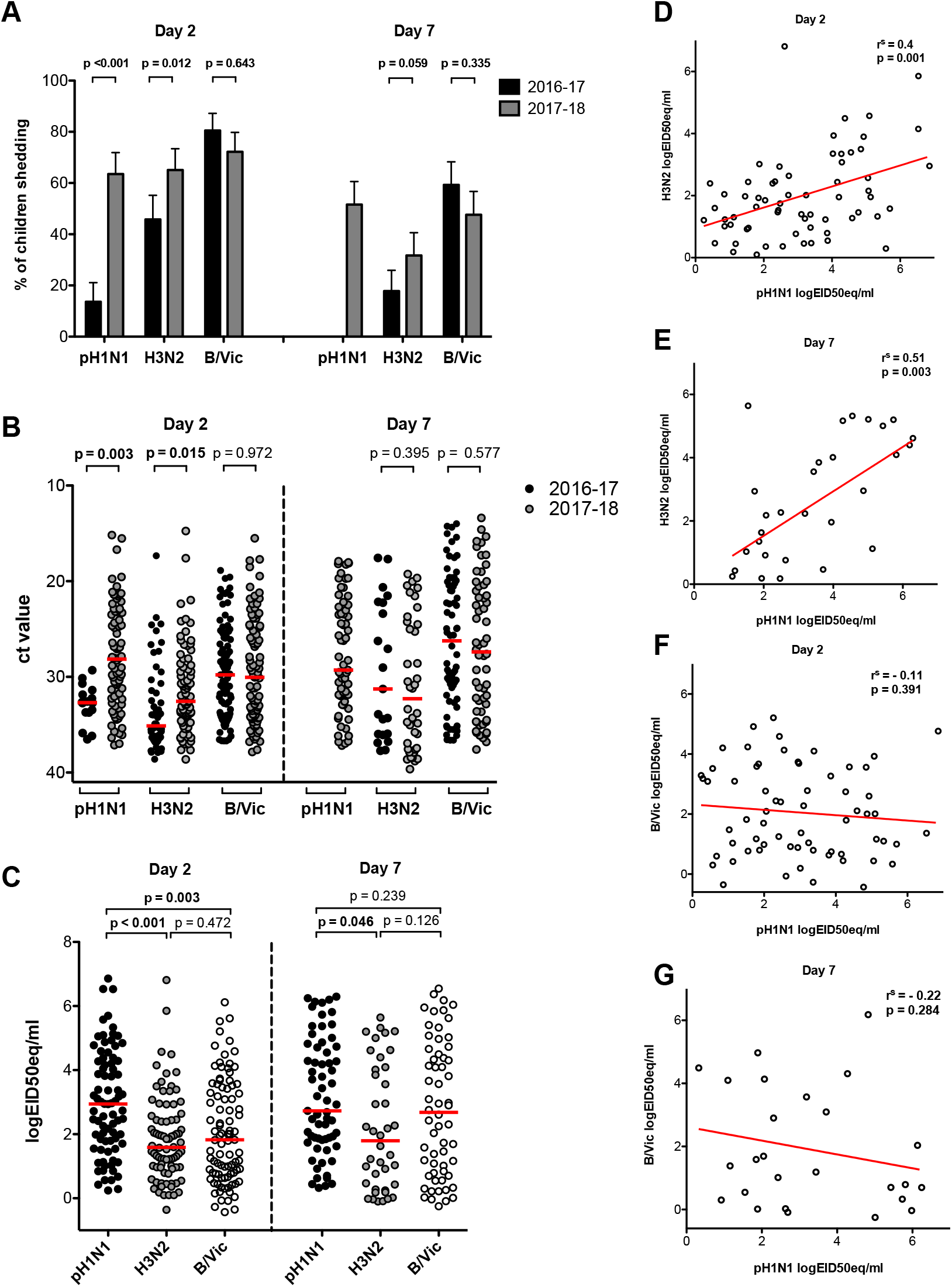
Shedding of strains in the nasopharynx following vaccination. **A**. Percentage of children shedding vaccine virus with 2016-17 formulation compared to the 2017-18 formulation, at day 2 and day 7. **B**. Cycle threshold (ct) values from reverse-transcriptase polymerase chain reaction (RT-PCR) for each strain, as a marker of viral load in the nasopharynx. Red bars indicate median ct values. Note lower ct values indicate higher viral loads **C**. Quantitative RT-PCR viral load (logEID50eq/ml = log 50% Egg Infectious Dose equivalents/ml) in children from the 2018 cohort for each strain. Red bars indicate median values. **D – G**. Correlation of nasopharyngeal viral loads in children given 2017-18 formulation between day 2 pH1N1 and H3N2 **(D)**, day 7 pH1N1 and H3N2 **(E)**, day 2 pH1N1 and B/Vic **(F)** and day 7 pH1N1 and B/Vic strains **(G)**. r^s^ = Spearman’s rank order coefficient. All displayed p values are Bonferroni-adjusted for multiplicity within each group of analyses.

To determine whether pre-existing adaptive immunity was responsible for poor Cal09 shedding, we calculated the predicted probability of shedding at each baseline HAI titre using logistic regression (figure 3A – D). While an inverse relationship was evident for H3N2 and B/Vic between baseline HAI titre and shedding, this was not evident Cal09, where low shedding is predicted even in seronegative children. In contrast, NY15 shedding was inversely related to the magnitude of the baseline HAI titre. Logistic regression also showed no associations between shedding and pre-vaccination T-cell response or HA-specific mucosal IgA for Cal09 (table S6). Similarly, no significant association between T-cell response or IgA and shedding was observed for H3N2 or B/Vic, after adjusting for baseline HAI titre (table S7). A lower nasopharyngeal viral load was seen at D2 and D7 for all three strains in baseline seropositive compared to seronegative children given the 2017-18 formulation (figure 3E), further emphasising the importance of serum antibody in this process.

**Figure 3.**
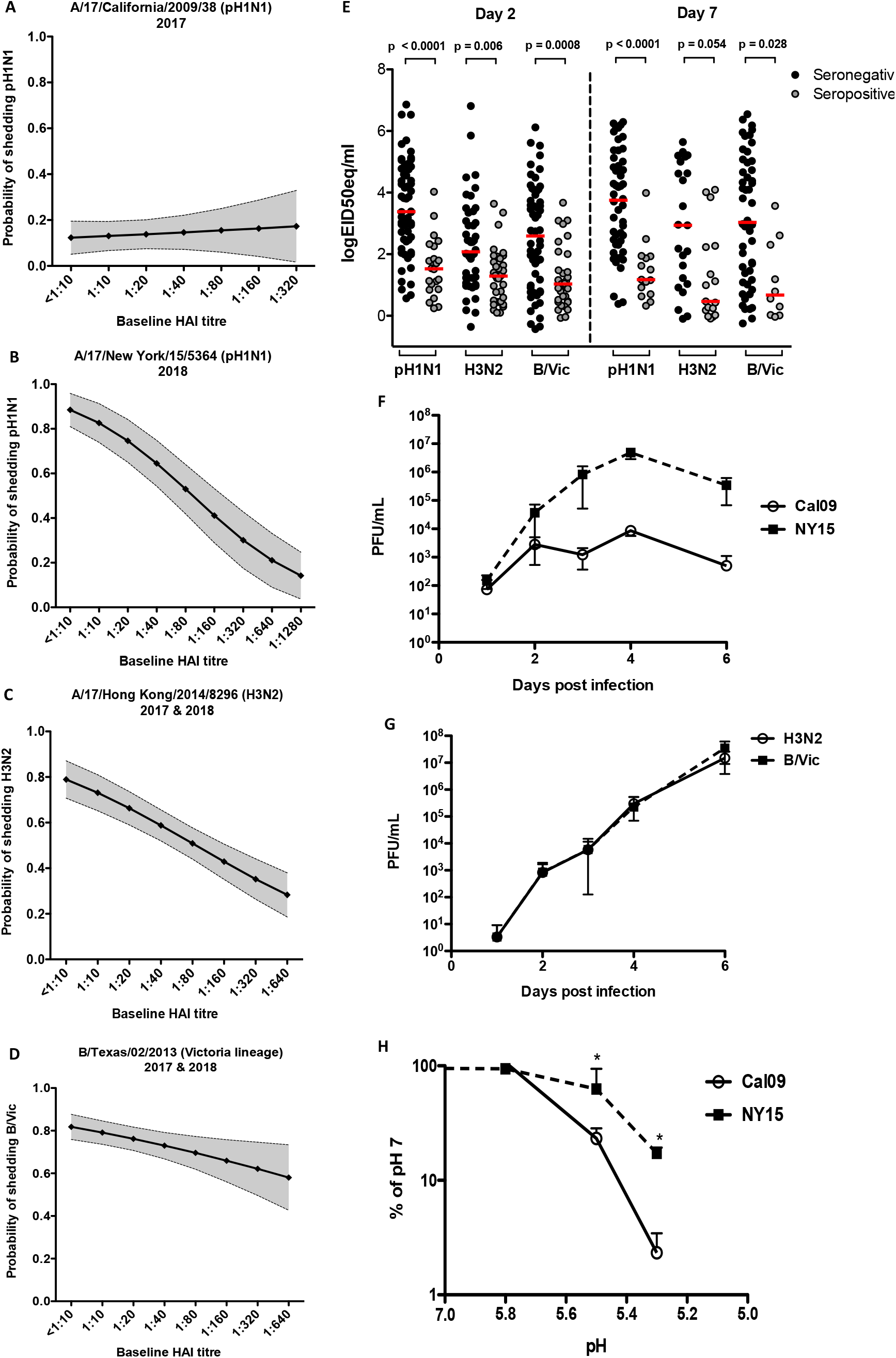
Impact of baseline serum antibody on LAIV strain shedding in the nasopharynx and replicative ability of viruses in primary epithelial cell cultures, where there is no influence of adaptive immunity. Predicted probability from logistic regression of vaccine strain shedding at day 2 post-LAIV at a given baseline serum haemagglutination inhibition (HAI) titre to each matched strain. Plotted are predicted mean proportions and 95% confidence intervals. Data shown for Cal09 pH1N1 **(A)**, NY15 pH1N1 **(B)**, H3N2 **(C)** and B/Victoria **(D)**. Upper limit is based on maximum observed HAI titre in the dataset. Where data from 2017 and 2018 were combined for H3N2 and B/Vic, results were adjusted for year (see table S8 for raw data). **(E)** Nasopharyngeal viral load (log10 50% Egg Infectious doses/ml equivalents, logEID50eq/ml) at day 2 and day 7 following the 2017-18 LAIV, with participants stratified by baseline serostatus to vaccine HA and NA-matched influenza strains. Replication of pH1N1 **(F)** or H3N2 and B/Victoria **(G)** vaccine strains in primary nasal epithelium. Effect of pH on vaccine strain growth in vitro **(H)**. Mean +/− standard deviation is shown. * p<0.05. PFU/ml = Plaque-forming Units/millilitre.

In seronegative children, shedding at D2 was seen in 12.7% (7.0-21.8) for Cal09 compared to 90.6% (81.0-95.6) for NY15 (p<0.001); in 75.0% (56.6-87.3) for H3N2 with 2016-17 LAIV compared to 82.1% (70.2-90.0) with 2017-18 LAIV (p=0.631); and in 83.9% (75.1-90.0) for B/Vic with 2016-17 LAIV compared to 79.2% (68.4-86.9) with 2017-18 LAIV (p=0.566). Ct value comparisons between 2016-17 and 2017-18 LAIV strains in seronegative children revealed the only significant difference as being a lower D2 ct (higher viral load) of NY15 compared to Cal09 (p<0.001, figure S5).

Monovalent vaccine strain replication was tested in primary hNECs cultured at airliquid interface, to see whether *in vitro* kinetics (in the absence of adaptive immune responses) reflected Cal09 and NY15 shedding in children. NY15 replication was greater than Cal09 (figure 3F), whereas H3N2 and B/Vic growth was equivalent and also superior to Cal09 (figure 3G). As stability in acidic environments in the upper respiratory tract (URT) may be important for replicative ability, we quantified Cal09 and NY15 after exposure to varying pH. This demonstrated greater stability of NY15 in acidic environments compared to Cal09 (figure 3H).

The 2016-17 LAIV resulted in significantly less children seroconverting to pH1N1 (5.1%, 1.9-10.7) compared to both H3N2 (22.0%, 14.9-30.6, p<0.001) and B/Vic (33.9%, 25.4-43.2, p <0.001). There was a significant increase in pH1N1 seroconversion with the 2017-18 LAIV (19.1%, 13.2-26.8, figure 4A), with no difference in H3N2 (27.8%, 20.7-36.2) or B/Vic (23.8%, 17.2-32.0). The improved seroconversion to pH1N1 with NY15 compared to Cal09 was especially evident in seronegative children (37.5%, 26.7-49.8, compared to 7.6%, 2.8-15.8, figure 4A), with a significant difference in HAI geometric mean fold rise for pH1N1 in 2017-18 (figure 4B).

**Figure 4.**
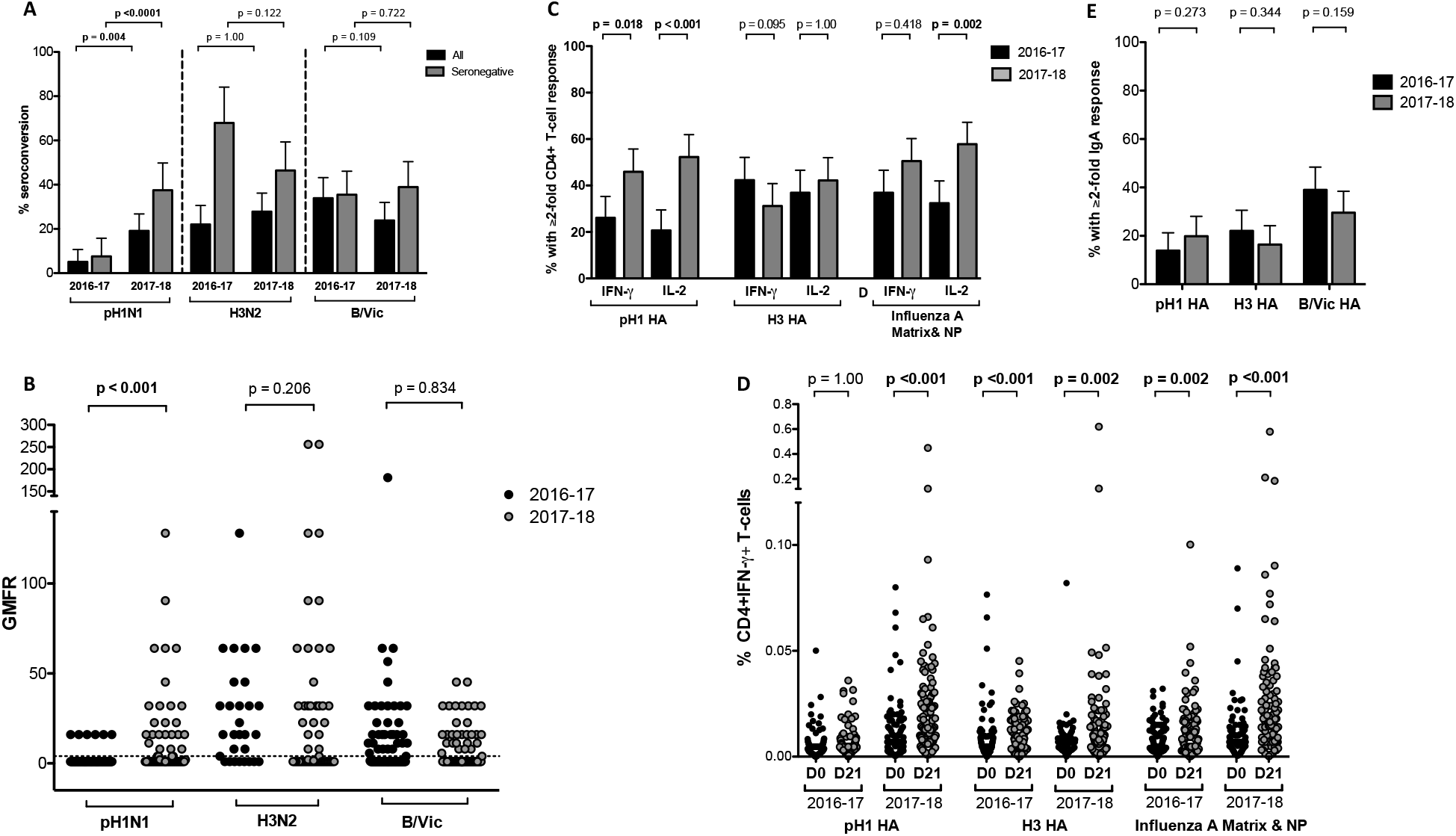
Improved immunogenicity to pH1N1 with the 2017-18 LAIV formulation. **(A)** Percentage of children seroconverting to each LAIV strain, comparing 2016-17 and 2017-18 formulations. **(B)** Geometric mean fold rise (GMFR) in serum haemagglutination inhibition titre from baseline to day 21, comparing children seronegative at baseline given 2016-17 and 2017-18 LAIVs. **(C) and (D)** Influenza-specific CD4+ T-cell responses to vaccine strain-matched pH1 haemagglutinin (HA; Cal09 or NY15 in respective years), H3 HA, influenza A matrix and nucleoprotein (both matched to LAIV backbone) peptide pools, comparing 2016-17 and 2017-18 LAIVs. **D.** Percentage of children with a 2-fold rise in influenza-specific CD4+ T-cell responses at day 21 after 2016-17 and 2017-18 LAIV. **(E)** Percentage of influenza-specific mucosal IgA responders given the 2016-17 and 2017-18 LAIVs. Displayed p values are Bonferroni-adjusted for multiplicity within each group of analyses. Error bars on plots displaying percentage of responders represent the upper 95% confidence interval. NP = nucleoprotein. IFN-γ = interferon gamma.

We were able to detect influenza-specific CD4+IFN-γ+ and/or CD4+IL-2+ and CD8+IFN-γ+ T-cell responses both at baseline and following vaccination. Although the magnitude of CD8+ responses was generally higher, LAIV-induced responses were predominantly CD4+ (figures 4C, 4D, S6, S7). The 2016-17 LAIV did not induce significant pH1 HA-specific CD4+IFN-γ+ or IL-2+ responses, whereas H3 HA- and A/MNP-specific responses were significantly increased from baseline (figures 4D and S6). In contrast, the 2017-18 LAIV induced significant pH1 HA-specific CD4+ T-cells at D21. Accordingly, the proportion of children with ≥2-fold rise in pH1 HA-specific CD4+ T-cell responses was significantly higher with the 2017-18 than 2016-17 LAIV: 45.9% (36.3-55.7) compared to 26.1% (18.2-35.3) for CD4+IFN-γ+ and 52.3% (42.5-61.9) compared to 20.7% (13.6-29.5) for CD4+IL-2+ responses (figure 4C). B/Vic HA- and influenza B MNP-specific CD4+ responses were also induced (figure S8). No significant change in the proportion of mono/dual-functional CD4+ T-cell responses was seen after vaccination (figure S9). The percentage of children with an influenza-specific mucosal IgA response to pH1N1 was not significantly different between the 2016-17 (13.9%, 8.0-21.3) and 2017-18 (19.8%, 13.1-28.1) LAIVs (figure 4E).

The impact of shedding on immunogenicity was explored using H3N2 data, as the largest sample size immunised with the same antigen and with T-cell data available. Seroconversion and T-cell responses were highest in children with shedding at both D2 and D7 (figures 5A and 5B). Multivariable logistic regression showed a significant impact of this prolonged shedding on the odds of seroconversion (figure 5D, table S9) and CD4+ T-cell response (figure 5E, table S10). No such relationship was seen with IgA responses (figures 5C and 5F, Table S11). Baseline HAI titre and induction of an H3 HA-specific CD4+IL-2+ response also increased the odds of seroconversion. Similar findings were observed in B/Vic and NY15 pH1N1 datasets, albeit with smaller sample sizes (tables S12 – S15).

**Figure 5.**
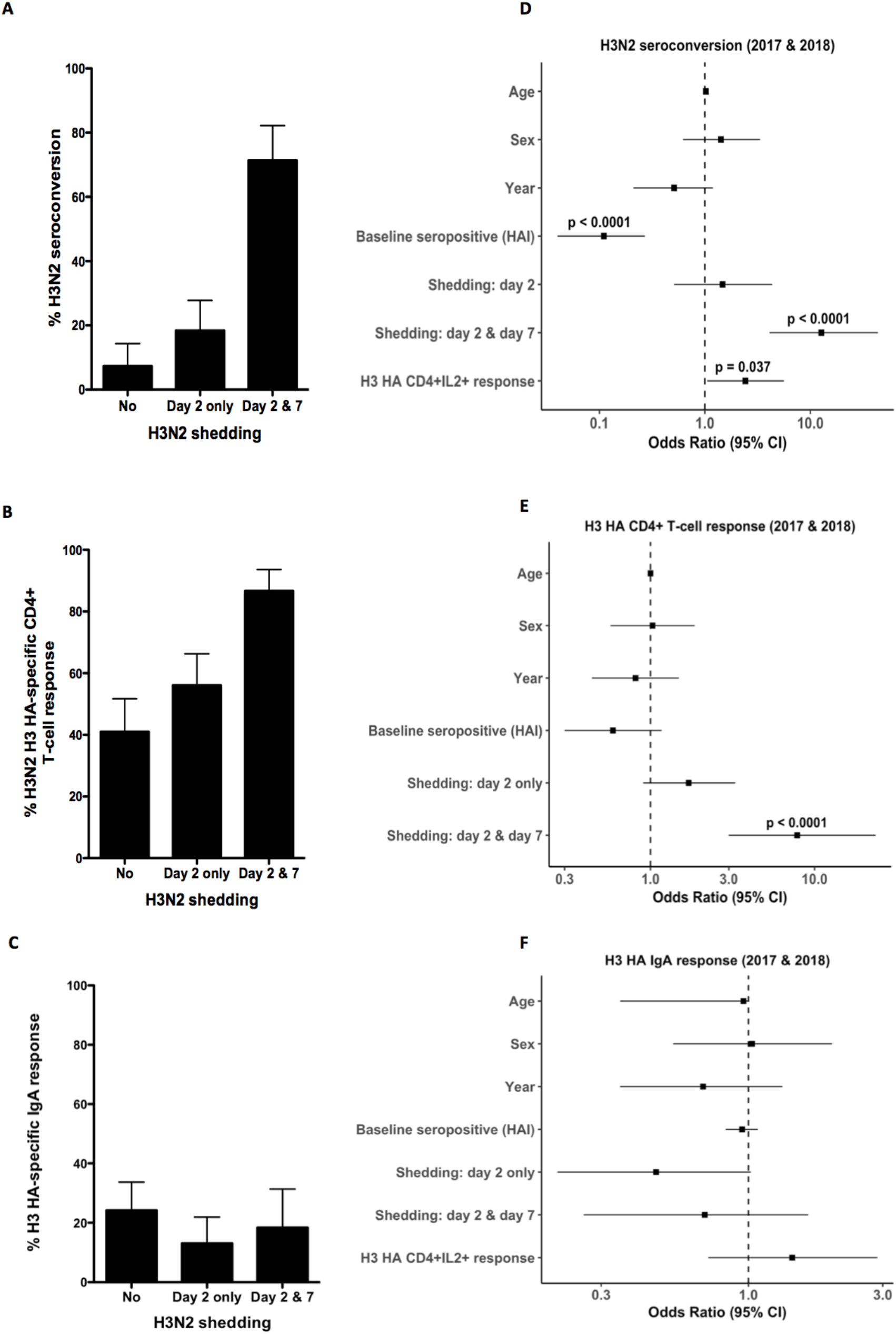
The impact of nasopharyngeal shedding on immune responses to LAIV. Percentage of children with **(A)** seroconversion, **(B)** 2-fold rise in haemagglutinin (HA)-specific CD4+ T-cell response (IFN-γ and/or IL-2) and **(C)** 2-fold rise in influenza HA-specific mucosal Immunoglobulin A (IgA) with each shedding category (no shedding, shedding at day 2 only, shedding at day 2 and day 7), using H3N2 data as an example (combined 2016-17 and 2017-18 data). Children without complete T-cell data (24/244) were excluded. The small number of children with shedding detected only at day 7 but not day 2 (12/244) were also excluded from the analysis. Odds ratio and 95% confidence intervals (CI) from multivariable logistic regression assessing the impact of shedding and other variables on **(D)** seroconversion, **(E)** CD4+ T-cell induction. **(F)** Odds ratio and 95% CI from univariate logistic regression assessing the impact of shedding on IgA response to H3N2. Odds ratio for shedding categories are calculated using no shedding as the reference.

## Discussion

We demonstrate limited shedding, *in vitro* Cal09 replication, and low immunogenicity following the 2016-17 LAIV in Gambian children, providing an explanation for the lack of efficacy observed in the RCT from neighbouring Senegal.^2^ Following the switch to NY15, a significant increase in replication was observed, along with improved serum humoral and cellular immunogenicity. No competitive inhibitory impact of enhanced pH1N1 replication was seen on H3N2 or B/Vic replication or immunogenicity. Our data also demonstrate that shedding for a longer duration is important for immunogenicity and that viral replicative fitness should be considered alongside antigenicity when selecting vaccine strains. These data represent the first reported LAIV immunogenicity data from African children to date. They make a case for further studies of LAIV efficacy in Africa, but are also of relevance to the use of LAIV in other settings.

A recent study of Ann-Arbor LAIV showed improved shedding and HAI seroconversion with an updated A/Slovenia/2015 pH1N1 strain.^20^ Parallel findings in two distinct paediatric cohorts, using two different LAIVs, provide strong support for Cal09 replicative fitness being culpable for recent problems. Our finding that limited Cal09 shedding is unlikely to be due to pre-existing immunity further supports this and argues against the notion that reduced LAIV effectiveness in the US may have been due to repeated vaccination in previous years.^21^

In an earlier study using the Ann-Arbor LAIV, pre-pandemic seasonal H1N1 shedding was found to be higher than for H3N2 or influenza B.^22^ Why pH1N1 Cal09 replication is impaired is uncertain. HA or NA residues must be responsible, as the remaining six viral gene segments in LAIV are consistent between Cal09, NY15 and H3N2 strains. Differences in Cal09 HA thermostability, sialic acid receptor binding and pH sensitivity are potential explanations for the lower replication observed.^1^ These phenotypes are important for optimal replication in the human URT. In particular, URT pH in children may be lower than for adults^23^ and have a deleterious impact on replication of viruses with labile HA. The pH1N1 virus first crossed into humans in 2009 and has subsequently circulated as a human seasonal virus. During this time, changes in HA stability and receptor binding properties may have adapted the virus to replicate better in the human URT.^24^ Thus, the more recent HA from A/Michigan/45/2015-like viruses of 2015 could have conferred enhanced shedding to LAIV pH1N1 components.

We were able to mirror our findings in a primary hNEC model. These cells display a mildy acidic apical surface environment akin to that seen in the human URT. This strategy may represent a practical method for evaluating vaccine virus replication prior to strain choice. As cell lines traditionally used to culture influenza viruses (e.g. MDCK) may not truly reflect replication in the URT, these subtleties were previously under-appreciated.^25^ Ultimately, a greater understanding of the viral genetic determinants of LAIV replicative fitness will be required to select optimal vaccine formulations.

Our study also emphasises the multifaceted nature of LAIV-induced immunity. While seroresponse (the traditional correlate of protection post-IIV) is modest, LAIV also induces mucosal IgA and T-cell responses. In our cohort, T-cell responses were elicited in a larger percentage of children than mucosal or serum antibodies, highlighting the importance of assessing cellular immunity in LAIV studies. Using the 2017-18 formulation, a CD4+ IFN-γ+ and/or IL-2+ T-cell response was seen in 55% - 68% of children to the influenza antigens tested, with approximately 80% showing a response to HA and/or MNP (figure S10). LAIV has been shown to provide protection in the absence of humoral immunity^26^ and T-cell-mediated immunity thought to play an important role.^27^

Unlike serum antibody and T-cell responses, we did not see a significant increase in mucosal IgA response with NY15. This is in keeping with findings after one dose of the updated Ann-Arbor LAIV, although a better response was seen after two doses.^20^ A recent immunogenicity study of Nasovac-S in Bangladesh also found that unlike serum antibody, nasal pH1N1-specific IgA was induced despite a lack of Cal09 shedding.^28^ The RCT these data were generated from reported LAIV efficacy of 50.0% (9.2–72.5) against pH1N1.^29^ Furthermore, in contrast to seroconversion and T-cell response, we found no significant association between shedding and IgA response. Taken together, these data suggest the mechanisms and requirements for serum antibody and mucosal IgA induction by LAIV may be distinct.

Our study has limitations. Participants were vaccinated with one LAIV dose, in keeping with the pre-qualification license from WHO and RCTs in Senegal and Bangladesh.^2,29^ Our findings may therefore not be generalisable to children in high income countries who receive booster doses and yearly influenza vaccination. We were also unable to confirm viral shedding with an LAIV-specific RT-PCR in all participants due to lower sensitivity compared with the HA-specific RT-PCR. However, by conducting the study outside of peak influenza season and baseline screening for influenza by RT-PCR, it is unlikely that our results were affected by wild-type influenza infections. Our shedding data at D2 are also similar to those reported from Senegal using Nasovac-S (Cal09:19.0%, H3N2:48.0% and influenza B:66.0%).^2^ As we measured shedding with RT-PCR and not culture, we are unable to confirm to what degree shedding reflected viable viruses. Finally, an important unanswered question from our study is whether NY15 and related pH1N1 strains will result in improved LAIV effectiveness. Data from the 2017-18 UK season estimates the vaccine effectiveness (Ann-Arbor LAIV) to be 90.3% against pH1N1 in 2-17-year-old children.^30^ However, owing to low-level circulation of pH1N1, the precision around this estimate was low (95% CI 16.4-98.9). Our findings suggest improved effectiveness can indeed be expected with the updated LAIV and if so, would support wider use of LAIV in the prevention of influenza.

## Acknowledgements

We gratefully acknowledge the study participants and parents who took part in the study. We also acknowledge the dedicated team of field and nursing staff lead by Janko Camara and Sulayman Bah, as well as Isatou Ndow for clinical trial organisation, the research support and clinical trials support offices at the MRC Unit The Gambia at LSHTM. We are most grateful to the Serum Institute of India Pvt Ltd for donating the vaccines used in this study. We acknowledge the help of Aminata Ngatou Vilane and Sheikh Jarju in establishing the RT-PCR assays. We are grateful to Dr. Yanchun Peng for help with overlapping peptide pools for influenza T-cell assays.

