## Supplementary information for "Use of a pandemic H1N1 strain with updated haemagglutinin and neuraminidase results in increased nasopharyngeal shedding and improved immunogenicity to Russian-backbone live attenuated influenza vaccine among children aged 2 – 4 years old: an open-label, prospective, observational, phase 4 study in"

### Table of Contents

Inclusion and Exclusion Criteria…………………………………………………………………......Page 2
Summary of parent study profile……………………………………………………………………..Page 3
- Figure S1
- Table S1

Details of T-cell assays using flow cytometry…………………………………………..………...Page 5
- Table S2
- Figure S2
- Figure S3
- Table S3

Details of RT-PCR assays used in shedding endpoints……………………………….…………...Page 7
- Table S4
- Figure S4
- Table S5

Associations between baseline immune responses and shedding of LAIV strains………………….Page 10
- Table S6
- Table S7
- Table S8
- Figure S5

Further details of T-cell responses to LAIV……………………………………………………...Page 12
- Figure S6
- Figure S7
- Figure S8
- Figure S9

Logistic regression analyses assessing the impact of shedding on immunogenicity to LAIV……...Page 14
- Table S9
- Table S10
- Table S11
- Table S12
- Table S13
- Table S14
- Table S15
- Table S16

Percentage of participants with T-cell responses to LAIV………………………….……………….Page 17
- Figure S10

References…………………………………………………………………………….………..Page 18

### Inclusion and Exclusion Criteria

#### *Inclusion Criteria*

- Healthy male or female child at least 24 months of age and less than 60 months of age at the time of study entry.
- Resident in the study area and with no plans to travel outside the study area during the period of subject participation.
- Informed consent for the study participation obtained from a parent (or guardian only if neither parent is alive or if guardianship has been legally transferred).
- Willingness and capacity to comply with the study protocol as judged by a member of the clinical trial team.

#### *Exclusion Criteria*

- Serious, active, medical condition, including but not limited to:
  - chronic disease of any body system
  - severe protein-energy malnutrition (weight-for-height Z-score of less than -3)
  - known genetic disorders, such as Down's syndrome or other cytogenetic disorder
- Active wheezing
- History of documented hypersensitivity to eggs or other components of the vaccine (including gelatin, sorbitol, lactalbumin and chicken protein), or with life-threatening reactions to previous influenza vaccinations.
- History of documented hypersensitivity to macrolide antibiotics
- History of Guillain-Barré syndrome.
- Receipt of aspirin therapy or aspirin-containing therapy within the two weeks before planned study vaccination.
- Any suspected or confirmed congenital or acquired state of immune deficiency including but not limited to primary immunodeficiencies including thymus disorders, HIV/AIDS, haematological or lymphoid malignancies.
- Any current immunosuppressive/immunomodulatory treatment or receipt of any such treatment within the six months preceding trial enrolment (for corticosteroids this is defined as a dose of prednisolone (or equivalent) of greater than 2mg/kg/day for one week or 1mg/kg/day for one month. The use of topical corticosteroids is not an exclusion criterion.
- The use of inhaled corticosteroids within the last one month.
- Receipt of an influenza vaccine within the past 12 months.
- Has any condition determined by investigator as likely to interfere with evaluation of the vaccine or be a significant potential health risk to the child or make it unlikely that the child would complete the study.
- Any significant signs or symptoms of an acute illness or infection including:
  - an axillary temperature of 38.0°C or above or documented fever of 38°C or above in the preceding 14 days.
  - Any acute respiratory infection within 14 days of enrolment visit.

**Summary of parent study profile (ClinicalTrials.gov identifier NCT02972957)**

The parent study is a phase 4, randomised controlled clinical vaccine trial: A Study of Intranasal Live Attenuated Influenza Vaccine Immunogenicity and Associations with the Nasopharyngeal Microbiome Among Children in The Gambia – The NASIMMUNE Study.

The sample size of the study and randomisation to LAIV vs unvaccinated arms was designed to document nasopharyngeal microbiome changes following LAIV compared to the control arm. Only participants receiving LAIV had samples taken for shedding and immunogenicity endpoints, therefore these represent data from a phase 4 observational study.

Children aged 24 – 59 months were recruited and randomised 1:1:1 to LAIV:LAIV:unvaccinated arms in both cohorts (target recruitment n = 110 in each group, total n = 330). This provided 90% power at alpha 0.05 to detect a 2-fold increase in density of relevant nasopharyngeal microbial taxa. The two LAIV groups followed identical study schedules other than the timing of an additional blood sample for exploratory immunological endpoints (at day 2 for one group and day 7 for the other), to minimise how many times each child was bled. In addition, in 2018, 35 children were recruited, given 1 dose of oral azithromycin, followed by the 2017-18 LAIV formulation 28 days later, for a pilot study to see if antibiotic therapy affected the interaction between LAIV and the nasopharyngeal microbiome. This group also had shedding and immunogenicity endpoints measured in an identical way to all other children given LAIV in the randomised groups. The summary study profile of the parent study is below (figure S1).

Data presented from the 2017-18 LAIV cohort therefore include data from n = 32 (of the total n = 126) who received azithromycin 28 days prior to LAIV. Shedding and immunogenicity endpoints from this group were not significantly different from those who did not received azithromycin (Table S1), therefore data are presented and analysed together.

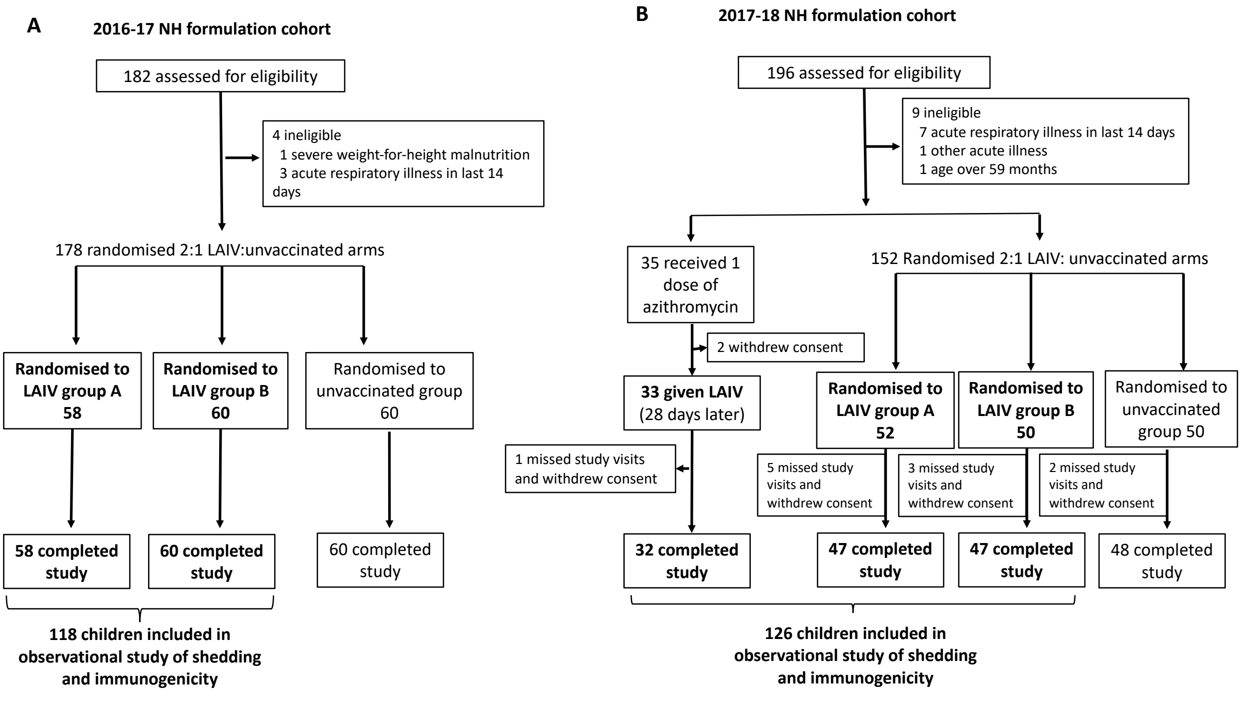

**Figure S1. Trial profile of parent study. Participants given LAIV are in bold and correspond to numbers presented in Figure 1.**

| Shedding and immune responses | 2017-18 cohort given azithromycin 28 days prior to LAIV (n = 32) % and 95% confidence interval | 2017-18 cohort: other (n = 94) % and 95% confidence interval | p value from Fisher's exact test (unadjusted) |
|---|---|---|---|
| Children shedding pH1N1 at day 2 | 65.62 (46.81-81.43) | 62.77 (52.18-72.52) | 0.8847 |
| Children shedding H3N2 at day 2 | 65.62(46.81-81.43) | 64.89 (54.36-74.46) | 1 |
| Children shedding B/Vic at day 2 | 78.12 (60.03-90.72) | 70.21 (59.9-79.21) | 0.4762 |
| Children shedding pH1N1 at day 7 | 46.88 (29.09-65.26) | 53.19 (42.61-63.56) | 0.7916 |
| Children shedding H3N2 at day 7 | 34.38 (18.57-53.19) | 30.85 (21.73-41.22) | 0.8529 |
| Children shedding B/Vic at day 7 | 50.0 (31.89-68.11) | 46.81 (36.44-57.39) | 0.8758 |
| Seroconversion pH1N1 | 15.62 (5.28-32.79) | 20.21 (12.63-29.75) | 0.7751 |
| Seroconversion H3N2 | 25.0 (11.46-43.4) | 28.72 (19.86-38.98) | 0.884 |
| Seroconversion B/Vic | 28.12 (13.75-46.75) | 22.34 (14.39-32.1) | 0.6507 |
| pH1 HA IgA response | 19.35 (7.45-37.47) | 19.54 (11.81-29.43) | 1 |
| H3 HA IgA response | 10.34 (2.19-27.35) | 18.89 (11.41-28.51) | 0.3602 |
| B/Vic HA IgA response | 25.81 (11.86-44.61) | 30.77 (21.51-41.32) | 0.7678 |
| CD4+IFN-gamma+ pH1 HA response | 46.43 (27.51-66.13) | 46.25 (35.03-57.76) | 1 |
| CD4+IFN-gamma+ H3 HA response | 14.29 (4.03-32.67) | 36.25 (25.79-47.76) | 0.05321 |
| CD4+IFN-gamma+ B/Vic HA response | 46.43 (27.51-66.13) | 31.25 (21.35-42.59) | 0.2234 |

**Table S1. Comparison of shedding and immunogenicity endpoints in individuals in the 2017-18 LAIV cohort given one dose of azithromycin 28 days prior to LAIV with those who did not receive any antibiotics.**

### Details of T-cell assays using flow cytometry

| Marker | Fluorochrome | Manufacturer | Clone |
|---|---|---|---|
| CD4 | PerCP-Cy5.5 | Biolegend | RPA-T4 |
| CD8 | FITC | Biolegend | RPA-T8 |
| IL-2 | PE | Biolegend | MQ1-17H12 |
| IFN-γ | APC | Biolegend | B27 |
| Viability dye | Live/dead® Aqua | Biolegend | NA |

**Table S2. Antibodies used in flow cytometry assays to detect influenza-specific T-cell response.** IFN-γ = interferon-gamma, IL-2 = interleukin-2.

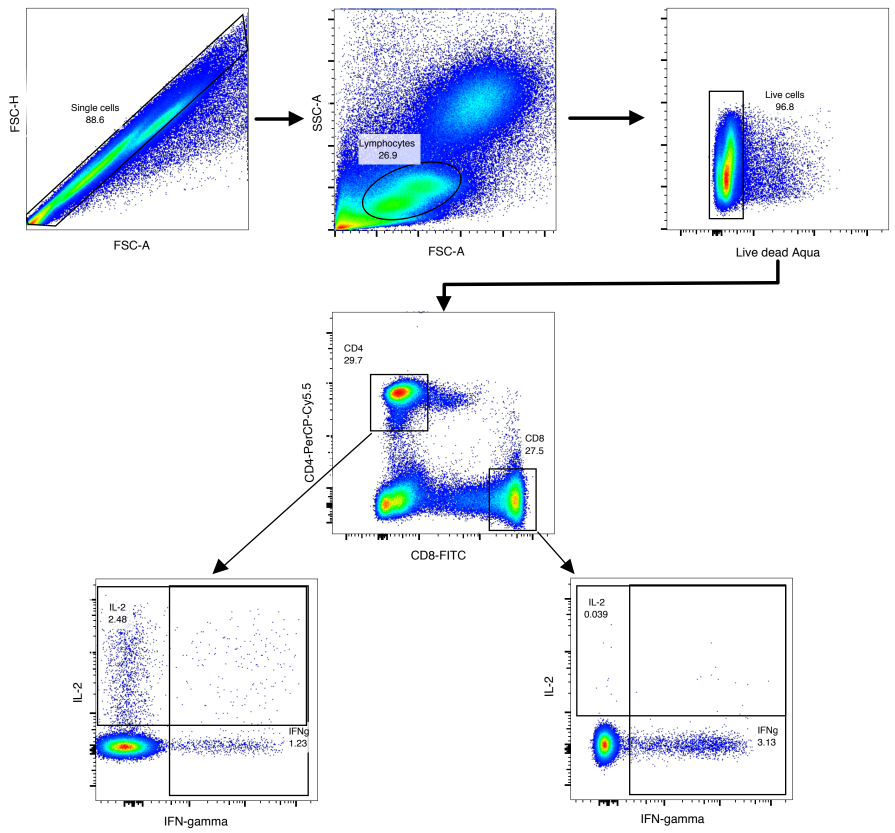

**Figure S2. Gating strategy for detection of IFN-γ and IL-2 in CD4+ and CD8+ T-cells. Displayed is from a sample following overnight stimulation with Staphylococcal Enterotoxin B (SEB).**

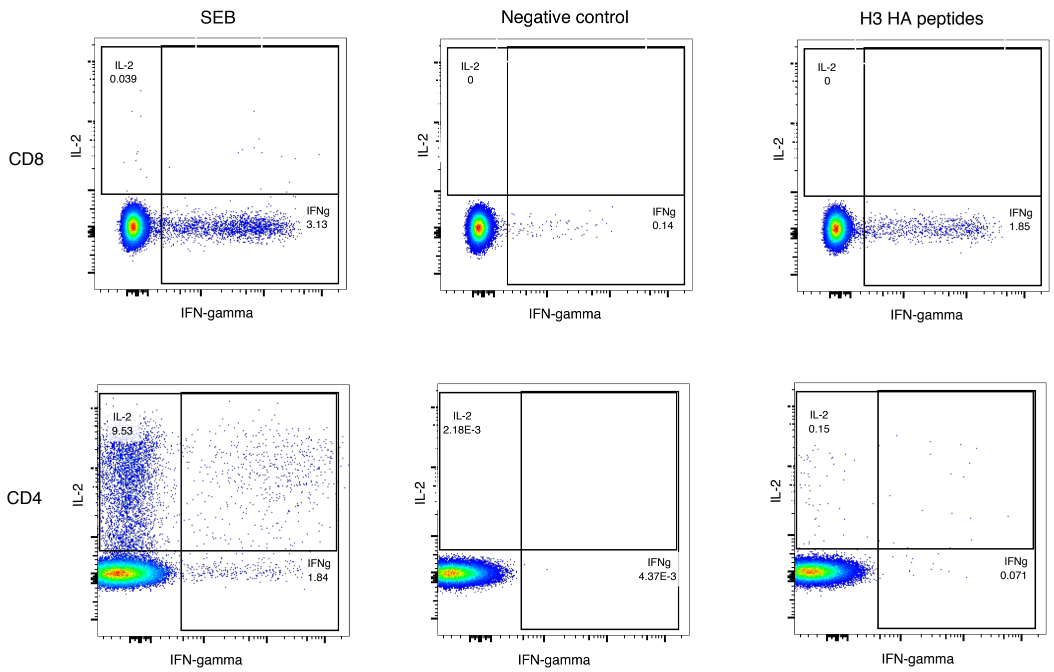

**Figure S3. CD8+ and CD4+ T-cell responses (IFN-γ and IL-2) following stimulation with Staphylococcal Enterotoxin B (SEB), negative control (co-stimulatory antibodies anti-CD28 and anti-CD48 alone) and an overlapping peptide pool matched to H3 haemagglutinin protein (A/H3N2 Hong Kong/4801/2014).**

| | 2017 data | 2018 data |
|---|---|---|
| **CD4 IFN-γ** | 0.009 | 0.014 |
| **CD4 IL-2+** | 0.015 | 0.016 |
| **CD8 IFN-γ+** | 0.047 | 0.110 |
| **CD8 IL-2+** | 0.005 | 0.014 |
| **CD4 IFN-γ+IL-2+** | 0.003 | 0.003 |
| **CD4 IFN-γ+IL-2-** | 0.007 | 0.013 |
| **CD4 IFN-γ-IL-2+** | 0.013 | 0.015 |

**Table S3.** Thresholds calculated based on the distribution of negative values after background subtraction, below which a positive value was also considered a null response.[1-3] IFN-γ = interferon-gamma, IL-2 = interleukin-2.

### Details of RT-PCR assays used in shedding endpoints

(i) RT-PCR conditions

Equine Arteritis Virus was added as an internal control followed by RNA extraction from NPS (260µl) using the QIAmp Viral RNA Mini Kit (Qiagen). Extracted RNA was reverse transcribed using 4X TaqMan™ Fast Virus 1-Step Master Mix (ThermoFisher) at 50˚C for 15 minutes, before denaturation at 95˚C for 60 seconds, followed by 50 cycles of amplification at 95˚C for 10 seconds and 60˚C for 30 seconds using a LightCycler® 96 (Roche). Cycle threshold (ct) values and positive sample calling were determined using automated LightCycler® software. Positive (seasonal influenza viruses) and negative (RNAse-free water) controls were included on each plate. All positive controls were consistently within two ct values.

| Name | Target | Label | Sequence |
|---|---|---|---|
| **HA-specific RT-PCR#** | | | |
| H1-Sw-1306F | pH1N1 HA | - | TGG ACT TAC AAT GCC GAA CT |
| H1-Sw-1423R | pH1N1 HA | - | CAG CCG TTT CCA ATT TCC TT |
| H1-sw-1357P2 | pH1N1 HA | TXR-BHQ2 | GAC TAC CAC GAT TCA AAT GTG AAG AAC T |
| H3-1541F1 | H3N2 HA | - | GAT GTR TAC AGA GAT GAA GCA TTA AAC A |
| H3-1600R | H3N2 HA | - | TAG GAT CCA ATC TTT GTA CCC TGA CTT |
| H3-1571P1 | H3N2 HA | YY-BHQ1 | AGC TCA ACT CCC TTG ATC TGG AAY CGG |
| INFB-HA-444F | Influenza B HA | - | ACC CTA CAR AMT TGG AAC YTC AGG |
| INFB-HA-524R | Influenza B HA | - | ACR GCC CAA GCC ATT GTT G |
| INFB-Vic499Pr | Influenza B Victoria lineage HA | Atto532-MGBFQ | ATC CGT TTC CAT TGG TAA |
| EAV-2043F | Equine Arteritis Virus (IC) | - | CTG TCG CTT GTG CTC AAT TTA C |
| EAV-2193R | Equine Arteritis Virus (IC) | - | AGC GTC CGA AGC ATC TC |
| EAV 2102P-2 | Equine Arteritis Virus (IC) | TXR-BHQ2 | TGC AGC TTA TGT TCC TTG CAC TGT GTT C |
| **Seasonal influenza internal gene RT-PCR#** | | | |
| INFAM-sense | Influenza A matrix | - | AAG ACC AAT CCT GTC ACC TCT GA |
| INFAM-sense3 | Influenza A matrix | - | AAG ACC AAT CTT GTC ACC TCT GA |
| INFAM-sense4 | Influenza A matrix | - | AAG ACC AAT TCT GTC ACC TCT GA |
| INFAM A-sense | Influenza A matrix | - | CAA AGC GTC TAC GCT GCA GTC C |
| INFAM- probe2-EDQ | Influenza A matrix | FAM-EDQ | TTT GTG TTC ACG CTC ACC GTG CC |
| INFB-NS779F | Influenza B NS gene | - | GTC TTA ATG AAG GAC ATT CAA AGC C |
| INFB-NS886R | Influenza B NS gene | - | TAA AGT TCT TCC GTG ACC AGT CTA |
| INFB 848P | Influenza B NS gene | YY-BHQ1 | GTC AAG AGC ACC GAT TAT CAC CAG AAG AG |
| **LAIV-specific internal gene RT-PCR*** | | | |
| MLenF427 | Influenza A | - | GCC TTT GGC CTG GTA TGT G |
| MLenR488 | Influenza A | - | CTA TGA GAC CTA TGC TGG GAG TCA |
| MLenP447 | Influenza A | FAM-BHQ1 | AAC CTG TGA ACA GAT TG |
| NSRussiaF377 | Influenza B NS gene | - | GAA AGT GCC TTG ATG ACA TA |
| NSRussiaR461 | Influenza B NS gene | - | TCT TTG TTG TTC ATG TCC CTC AAT |
| NSRussiaP426r | Influenza B NS gene | Atto532-BHQ1 | TGG GTC ATC AAC ATT TTC CGG TTC |

**Table S4. Primer and probe sequences used in reverse-transcriptase polymerase chain reaction (RT-PCR) assays.** #Routinely used in surveillance of influenza at the National Institute for Public Health and the Environment, Bilthoven, the Netherlands, which is one of the two locations of the National Influenza Centre in which these tests are performed under ISO 15189 accreditation for medical laboratories.[4] *Influenza A assays for LAIV-specific RT-PCR were based on previously published primers and probes that distinguish seasonal influenza A from LAIV influenza A master donor virus.[5] Influenza B assays for LAIV-specific RT-PCR were modified from previously published primers and probes for seasonal influenza B and incorporated key changes to match the B/Russia/69 sequence (which represents a close match the

Nasovac-S influenza B master donor virus) and differentiate from current circulating strains. IC – internal control. HA = haemagglutinin.

(ii) <u>Details of quantitative RT-PCR assay used with the 2017-18 cohort:</u>

A standard curve was included on each plate with two replicates of LAIV dilutions from $1{:}10^3$ to $1{:}10^6$ , with known log10 50% Egg Infectious Doses/ml (EID/ml). Testing during assay validation showed continued linearity up to a 1:10 dilution. Median RT-PCR efficiency (based on standard curve) was 98% (range 88-110%). Median standard curve $R^2$ was 0.998 (range 0.977-1). 95% LOD was determined using a serial dilution (10 replicates) of 2017-18 formulation vaccine (figure S3). LOD for pH1N1, H3N2 and Influenza B PCRs were 1.11 (95% CI 0.89-1.22), 0.91 (0.61-1.09) and 0.73 (0.5-0.85) EID/ml respectively (Figure S4 below).

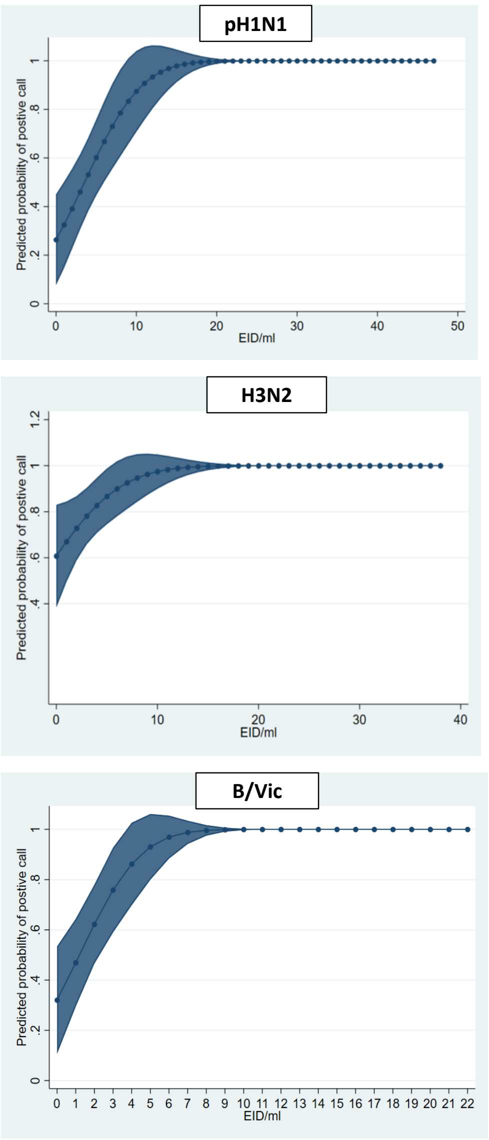

**Figure S4. 95% limit of detection (LOD) for pH1N1, H3N2 and B/Vic RT-PCR using HA-specific primer and probes sets.** Calculated using a 2-fold serial dilution of 10 LAIV replicates from $1{:}4 \times 10^6$ to $1{:}6.4 \times 10^7$ (after an initial 10-fold three-replicate dilution series from $1{:}10^3$ to $1{:}10^9$ to estimate the LOD for each RT-PCR). Displayed are predicted probability (and 95% confidence interval) of a positive reaction at each log10 50% egg infectious dose/ml (EID/ml) calculated using Stata 12.1 (Probit model).[6] LOD for pH1N1, H3N2 and Influenza B PCRs were 1.11 (95% CI 0.89-1.22), 0.91 (0.61-1.09) and 0.73 (0.5-0.85) EID/ml respectively.

(iii) LAIV-specific RT-PCR:

A 10-fold dilution series ($1:10^2 – 1:10^8$) of seasonal influenza strains and LAIV were tested with seasonal influenza and LAIV-specific sets (table S4). LAIV-specific assays detected LAIV but not seasonal influenza strains (pH1N1, H3N2 and B/Victoria), whereas as expected, seasonal influenza assays detected both. Despite optimisation of assay conditions, the maximum LAIV dilution detected by LAIV sets was at least one log10 lower than that detected by HA-specific sets (Table S5).

| Primer/probe set (target) | Highest LAIV dilution detected* |
|---|---|
| LAIV influenza A (matrix) | $1:10^6$ |
| LAIV influenza B (NS) | $1:10^5$ |
| Seasonal influenza A (matrix) | $1:10^7$ |
| Seasonal influenza B (NS) | $1:10^6$ |
| pH1N1 (HA) | $1:10^7$ |
| H3N2 (HA) | $1:10^7$ |
| B/Vic (HA) | $1:10^8$ |

**Table S5. Ability of LAIV and seasonal influenza primer/probe sets to detect LAIV strains.** *results from 10-fold dilution series from $1:10^2$ to $1:10^8$ run in quadruplicate. Displayed are highest dilutions at which at least one positive reaction was seen. Haemagglutinin (HA)-specific sets appeared to be more sensitive than LAIV (matrix and NS1)-specific sets. Of note the influenza A matrix sets should amplify influenza A LAIV master donor virus from both pH1N1 and H3N2 vaccine strains, thus approximately double the amount of virus should be available as a target compared to influenza B internal gene sets or any HA-specific sets.

**Associations between baseline immune responses and shedding of LAIV strains**

| | | Univariate | |
|---|---|---|---|
| | | OR | p value |
| Age (in months) | | 0.99 | 0.779 |
| Sex | | 1.08 | 0.884 |
| Z-score (weight-for-height) | | 1.32 | 0.343 |
| Baseline seropositivity (HAI titre) | | 1.25 | 0.684 |
| Baseline IgA (% of total IgA)* | 2nd tertile | 3.33 | 0.276 |
| | 3rd tertile | 2.06 | 0.515 |
| Baseline CD4+ T-cell MNP IFN-γ response | | 0.27 | 0.214 |
| Baseline CD8+ T-cell MNP IFN-γ response | | 0.31 | 0.277 |

**Table S6. No association between baseline immunity and shedding at day 2 of 2016-17 LAIV pH1N1 (Cal09) strain in univariate logistic regression.** OR = odds ratio. HAI = haemagglutination inhibition. MNP = matrix and nucleoprotein. IFN-γ = interferon gamma. *1st tertile used as reference for calculating ORs.

| Baseline immune response | | Strain | Univariate | | Adjusted for baseline HAI titre | |
|---|---|---|---|---|---|---|
| | | | OR | p value | OR | p value |
| **Influenza-specific IgA (% of total)** | 2nd tertile | pH1N1 NY15 | 0.6 | 0.250 | - | - |
| | 3rd tertile | | 0.47 | 0.131 | - | - |
| **Influenza-specific IgA (% of total)†** | 2nd tertile | H3N2 | 0.63 | 0.169 | 0.99 | 0.988†† |
| | 3rd tertile | | **0.31** | **<0.001** | 0.68 | 0.311†† |
| **Influenza-specific IgA (% of total)†** | 2nd tertile | B/Vic | 0.69 | 0.340 | - | - |
| | 3rd tertile | | 0.60 | 0.178 | - | - |
| **CD4+ T-cell A/MNP IFN-γ response** | | pH1N1 NY15 | 0.39 | 0.07 | - | - |
| **CD8+ T-cell A/MNP IFN-γ response** | | pH1N1 NY15 | 0.23 | 0.24 | - | - |
| **CD4+ T-cell A/MNP IFN-γ response†** | | H3N2 | **0.40** | **0.007** | 0.52 | 0.069†† |
| **CD8+ T-cell A/MNP IFN-γ response†** | | H3N2 | 0.75 | 0.43 | - | - |
| **CD4+ T-cell B/MNP IFN-γ response*** | | B/Vic | 0.33 | 0.082 | - | - |
| **CD8+ T-cell B/MNP IFN-γ response*** | | B/Vic | No baseline responses seen | | | |

**Table S7. Impact of pre-existing IgA and T-cell immunity on probability of shedding following LAIV.** †Combined 2017 & 2018 data. ††Year included in multivariable model. 1st tertile used as reference in IgA analyses. *Influenza B T-cell responses only available from 2018 cohort. No CD8+ T-cell responses to vaccine antigen-matched influenza B master donor virus matrix or nucleoprotein seen (above calculated thresholds defining a true response) in baseline samples. OR = odds ratio. MNP = matrix and nucleoprotein. HAI = haemagglutination inhibition. IFN-γ = interferon gamma.

| Baseline (reciprocal) HAI titre | pH1N1 2016-17 (Cal09) | pH1N1 2017-18 (NY15) | H3N2 2016-17 | H3N2 2017-18 | B/Vic 2016-17 | B/Vic 2017-18 |
|---|---|---|---|---|---|---|
| <10 | 10/79 | 58/64 | 21/28 | 46/56 | 78/93 | 57/72 |
| 7.1 | - | - | - | - | - | 1/1 |
| 10 | - | - | - | 1/1 | - | - |
| 20 | - | 0/1 | - | - | - | - |
| 28.3 | - | - | 1/2 | - | - | - |
| 40 | 0/1 | 1/4 | 3/4 | 7/8 | 2/2 | 1/2 |
| 56.6 | - | - | 1/3 | - | 1/1 | 0/1 |
| 80 | 1/14 | 4/9 | 7/22 | 5/12 | 6/7 | 2/4 |
| 113.1 | 1/3 | - | 2/3 | 0/1 | - | - |
| 160 | 3/15 | 7/15 | 14/34 | 7/20 | 5/7 | 12/16 |
| 226.3 | 0/2 | 3/4 | 1/3 | 3/3 | 1/2 | 5/9 |
| 320 | 1/4 | 4/17 | 2/13 | 10/16 | 2/5 | 7/11 |
| 452.6 | - | 0/3 | 0/1 | 1/2 | 0/1 | 1/2 |
| 640 | - | 3/8 | 1/4 | 2/7 | - | 4/7 |
| 905.1 | - | - | 1/1 | - | - | 1/1 |
| 1280 | - | 0/1 | - | - | - | - |

**Table S8. Number of children shedding LAIV strains at day 2 post-LAIV.** Denominator in each cell is the number of participants with the respective Haemagglutination inhibition (HAI) titre at baseline.

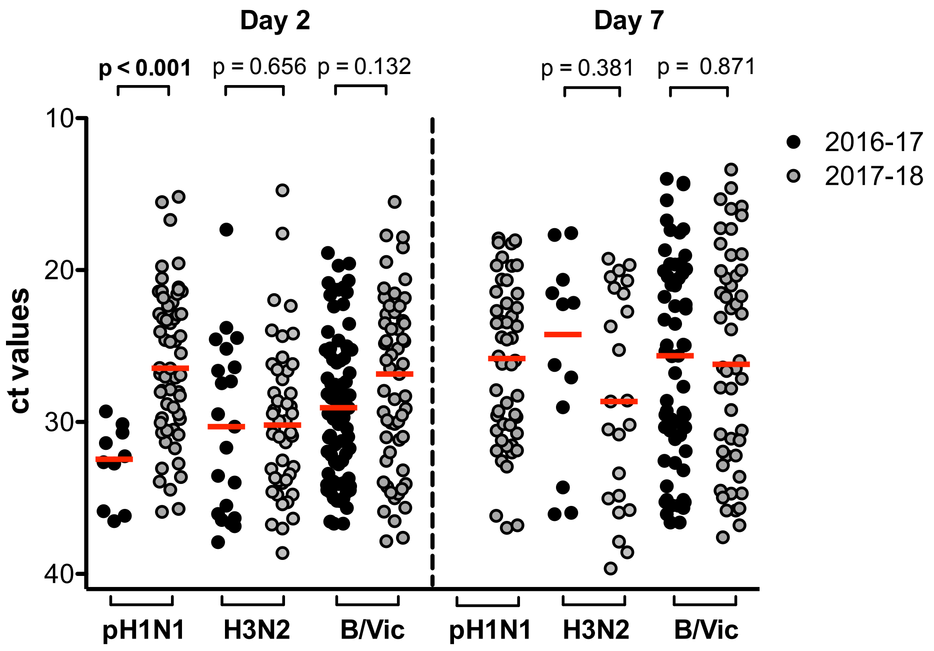

**Figure S5. Cycle threshold (ct) values from reverse-transcriptase polymerase chain reaction for each strain, as a marker of viral load in the nasopharynx, in children seronegative at baseline.** Red bars indicate median ct values. Note lower ct values indicate higher viral loads. All displayed p values are Bonferroni-adjusted for multiplicity within each group of analyses. No comparison of day 7 pH1N1 shedding possible as no pH1N1 was detected in samples at day 7 with the 2016-17 LAIV.

### Further details of T-cell responses to LAIV

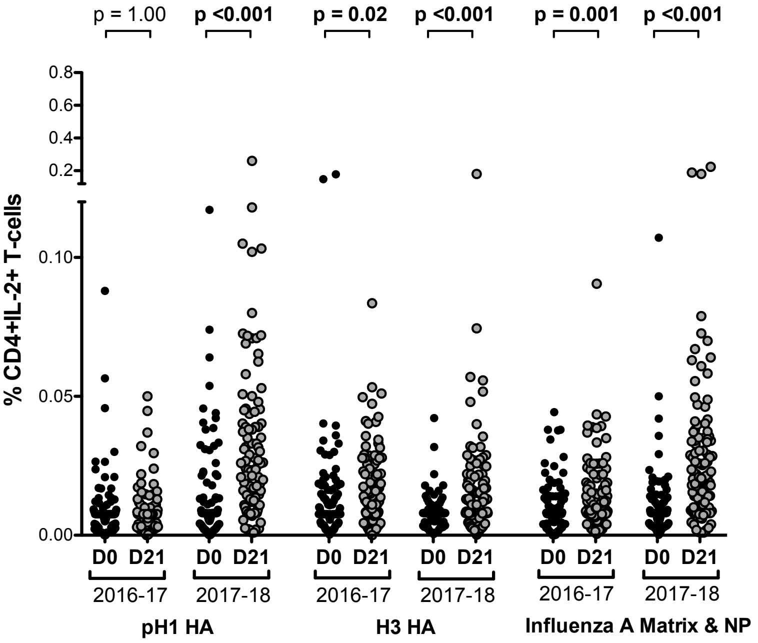

**Figure S6. CD4+IL2+ T-cell responses to influenza A antigens using the 2016-17 and 2017-18 LAIV formulations.** Significant increase from day 0 (D0) to day 21 (D21) of IL-2+ CD4+ T-cell responses to pH1 haemagglutinin (HA) seen only using 2017-18 formulation. CD4+IL-2+ T-cell responses to H3 HA, as well as matrix and nucleoprotein (NP) antigen seen using both formulations. IFN-γ = interferon gamma. IL-2 = interleukin 2. P values are Bonferroni-adjusted for multiple comparisons.

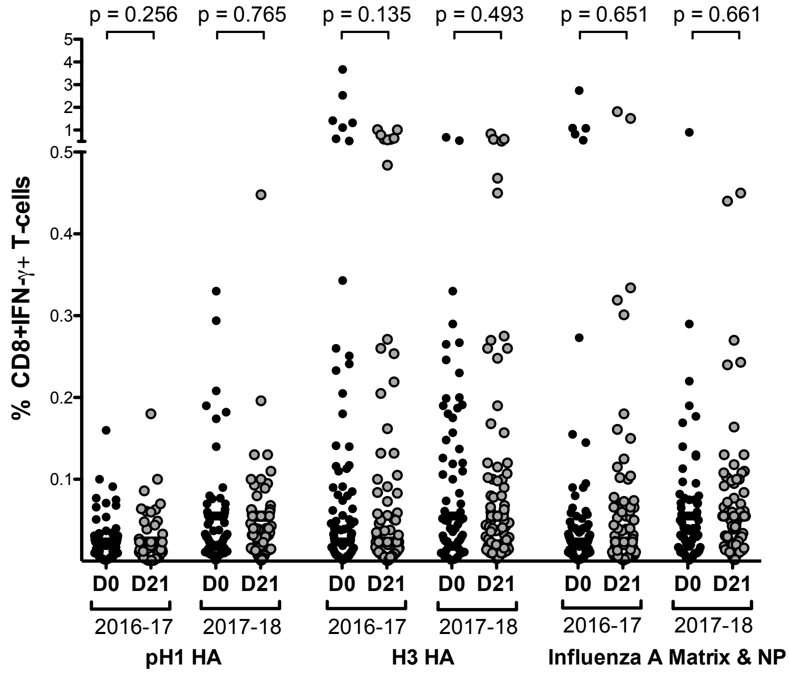

**Figure S7. CD8+ IFN-γ+ T-cell responses to influenza A antigens using the 2016-17 and 2017-18 LAIV formulations.** No significant increase from day 0 (D0) to day 21 (D21) seen to pH1 haemagglutinin (HA), H3 HA or matrix and nucleoprotein (NP) antigens seen using either formulation. IFN-γ = interferon gamma.

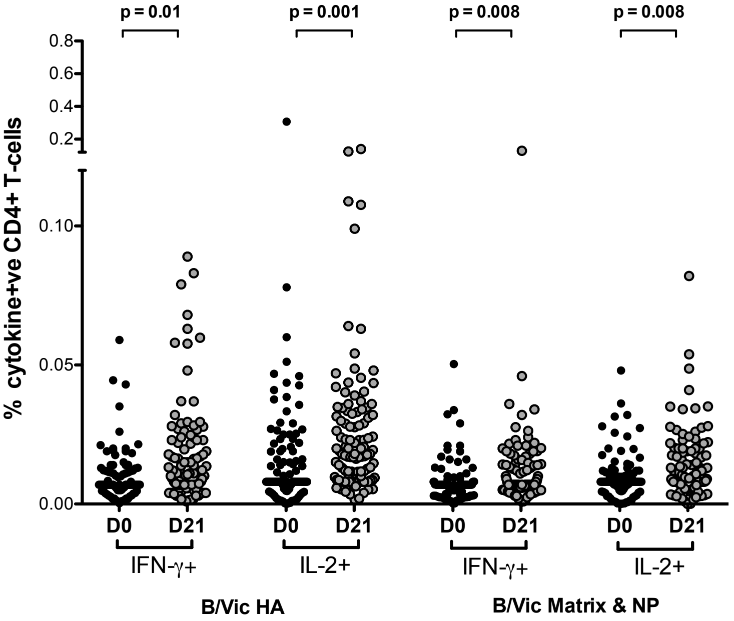

**Figure S8. Induction of CD4+ T-cell responses to influenza B antigens using the 2017-18 LAIV formulation.** Significant increase from day 0 (D0) to day 21 (D21) of IFN-γ+ and IL-2+ CD4+ T-cell responses to influenza B Victoria lineage (B/Vic) haemagglutinin (HA), as well as matrix and nucleoprotein (NP) antigen from Russian-backbone LAIV. IFN-γ = interferon gamma. IL-2 = interleukin 2. P values are Bonferroni-adjusted for multiple comparisons.

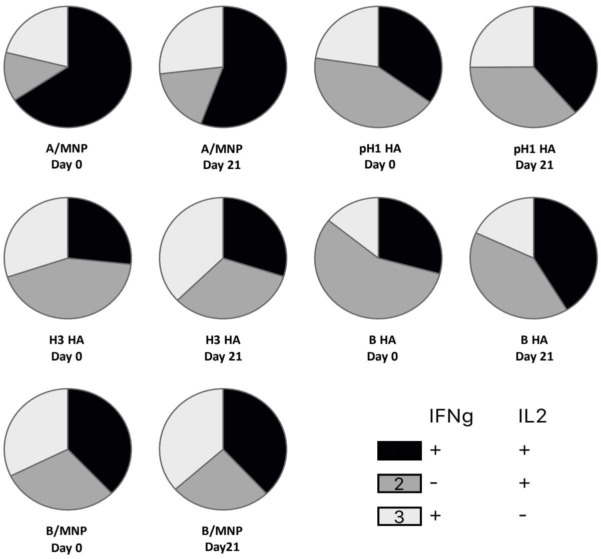

**Figure S9. The proportion of mono- and dual-functional CD4+ T-cell responses to influenza antigens tested**. No difference seen between baseline and day 21. Influenza B HA and matrix and nucleoprotein (MNP) data are from 2018 only, whereas others are combined 2017 and 2018 data. Significance between proportions of mono- and dual- functional responses across timepoints tested using the Permutation test (SPICE V6.0). IFNg = interferon gamma. IL-2 = interleukin 2.

### Logistic regression analyses assessing the impact of shedding on immunogenicity to LAIV

| H3N2 seroconversion (2017 & 2018) | | Univariate | | | Multivariable | | |
|---|---|---|---|---|---|---|---|
| | | OR | 95% CI | p value | OR | 95% CI | p value |
| Year | | 1.37 | 0.75 - 2.53 | 0.315 | 0.51 | 0.21 - 1.19 | 0.128 |
| Age | | 0.99 | 0.95 - 1.02 | 0.491 | 1.02 | 0.98 - 1.08 | 0.239 |
| Sex | | 1.15 | 0.63 - 2.13 | 0.648 | 1.42 | 0.62 - 3.33 | 0.406 |
| Baseline seropositive (HAI titre) | | **0.1** | **0.05 - 0.2** | **<0.0001** | **0.11** | **0.04 - 0.27** | **<0.0001** |
| Shedding | Day 2 only | 2.43 | 0.96 - 6.7 | 0.068 | 1.45 | 0.51 - 4.34 | 0.494 |
| | Day 2 and Day 7 | **26.7** | **10.29 - 78.52** | **<0.0001** | **12.69** | **4.1 - 43.6** | **<0.0001** |
| H3 HA CD4+IFN-γ+ response | | **1.71** | **0.92 - 3.19** | **0.086** | - | - | - |
| H3 HA CD4+IL-2+ response | | **4.95** | **2.61 - 9.7** | **<0.0001** | **2.42** | **1.05 - 5.62** | **0.037** |

**Table S9. Impact of shedding on seroconversion to H3N2.** Univariate and multivariable logistic regression using n = 210 children. Excludes 24/244 participants with missing T-cell data and 12/224 with shedding only detected at day 7 (2 children had both T-cell data missing and shedding only at day 7). Shedding coded as one variable with three levels (no shedding, day 2 only shedding, day 2 and day 7 shedding). H3 Haemagglutinin (HA) CD4+IL-2+ response taken forward to multivariable analysis due to overlap between this and the CD4+IFN-γ response. No interactions observed between variables included in the multivariable analysis. OR = odds ratio, CI = confidence interval, HAI = haemagglutination inhibition

| H3 HA CD4+IFN-γ and/or IL2 response (2017 & 2018) | | Univariate | | | Multivariable | | |
|---|---|---|---|---|---|---|---|
| | | OR | 95% CI | p value | OR | 95% CI | p value |
| Year | | 1.21 | 0.71 - 2.07 | 0.486 | 0.81 | 0.44 - 1.49 | 0.500 |
| Age | | 0.99 | 0.96 - 1.02 | 0.533 | 1.00 | 0.97 - 1.03 | 0.947 |
| Sex | | 1.05 | 0.61 - 1.79 | 0.866 | 1.03 | 0.57 - 1.86 | 0.915 |
| Baseline seropositive (HAI) | | **0.39** | **0.21 - 0.69** | **0.002** | 0.59 | 0.3 - 1.17 | 0.134 |
| Shedding | Day 2 only | 1.84 | 1.00 - 3.44 | 0.053 | 1.71 | 0.9 - 3.29 | 0.102 |
| | Day 2 and Day 7 | 9.38 | 3.8 - 26.85 | **<0.0001** | **1.56** | **2.99 - 23.5** | **<0.0001** |

**Table S10. Impact of H3N2 shedding on H3 HA-specific CD4+ T-cell response (IFN-γ and/or IL-2).** Univariate and multivariable logistic regression using n = 210 children. Excludes 24/244 participants with missing T-cell data and 12/224 with shedding only detected at day 7 (2 children had both T-cell data missing and shedding only at day 7). Shedding coded as one variable with three levels (no shedding, day 2 only shedding and day 2 and day 7 shedding). No interactions observed between variables included in the multivariable analysis. OR = odds ratio, CI = confidence interval, HAI = haemagglutination inhibition

<table>
<tr><th colspan="2" rowspan="2">H3 HA IgA response (2017 & 2018)</th><th colspan="3">Univariate</th></tr>
<tr><th>OR</th><th>95% CI</th><th>p value</th></tr>
<tr><td colspan="2">Year</td><td>0.69</td><td>0.35 - 1.32</td><td>0.268</td></tr>
<tr><td colspan="2">Age</td><td>0.96</td><td>0.93 - 1.00</td><td>0.071</td></tr>
<tr><td colspan="2">Sex</td><td>1.03</td><td>0.54 - 1.98</td><td>0.928</td></tr>
<tr><td colspan="2">Baseline seropositive (HAI)</td><td>0.95</td><td>0.83 - 1.08</td><td>0.413</td></tr>
<tr><td rowspan="2">Shedding</td><td>Day 2</td><td>0.47</td><td>0.21 - 1.02</td><td>0.062</td></tr>
<tr><td>Day 2 and Day 7</td><td>0.70</td><td>0.26 - 1.63</td><td>0.426</td></tr>
<tr><td colspan="2">H3 HA CD4+IFNg+ response</td><td>1.02</td><td>0.50 - 2.06</td><td>0.946</td></tr>
<tr><td colspan="2">H3 HA CD4+IL2+ response</td><td>1.43</td><td>0.72 - 2.87</td><td>0.304</td></tr>
</table>

**Table S11. Impact of H3N2 shedding on H3 HA-specific mucosal IgA response.** Univariate logistic regression using n = 228 children. Excludes 4/244 participants with missing IgA data and 12/244 with shedding seen only at day 7. Univariate analysis with T-cell independent variables done with n = 206 children (excludes a further 22 children with missing T-cell data). Shedding coded as one variable with three levels (no shedding, day 2 only shedding, day 2 and day 7 shedding). No interactions observed between variables included in the multivariable analysis. OR = odds ratio, CI = confidence interval, HAI = haemagglutination inhibition

| B/Vic seroconversion (2018) | Univariate | | | Multivariable | | |
|---|---|---|---|---|---|---|
| | OR | 95% CI | p value | OR | 95% CI | p value |
| Age | 0.95 | 0.90 - 1.00 | 0.06 | 0.96 | 0.89 - 5.16 | 0.26 |
| Sex | 1.96 | 0.81 - 5.03 | 0.15 | 1.75 | 0.58 - 5.56 | 0.32 |
| Baseline seropositive (HAI) | **0.06** | **0.01 - 0.21** | **0.0002** | **0.13** | **0.02 - 0.57** | **0.02** |
| Day 2 shedding | **10.07** | **1.95 - 184.9** | **0.03** | - | - | - |
| Day 7 shedding | **23.31** | **6.35 - 151.3** | **<0.0001** | **10.47** | **2.35 - 75.9** | **0.006** |
| B/Vic HA CD4+IFNg-γ+ response | 1.94 | 0.80 - 4.70 | 0.139 | - | - | - |
| B/Vic HA CD4+IL2+ response | **3.99** | **1.63 - 10.38** | **0.003** | 1.17 | 0.35 - 3.84 | 0.793 |

**Table S12. Impact of B/Vic shedding on B/Vic seroconversion.** Univariate and multivariable logistic regression using n = 109 children. Only participants from 2018 used as T-cell data were only generated to influenza B antigens in 2018. Excludes 17/126 participants with missing T-cell data. Day 2 and day 7 shedding variables included separately as combining into one variable (no shedding, day 2 only, day 2 and day 7) resulted in zero cell values (no seroconverters among those with no shedding). Only day 7 shedding taken forward to multivariable analysis. B/Vic Haemagglutinin (HA) CD4+IL-2+ response taken forward to multivariable analysis due to overlap between this and the CD4+IFN-γ response. No interactions observed between variables included in the multivariable analysis. OR = odds ratio, CI = confidence interval, HAI = haemagglutination inhibition.

| B/Vic HA CD4+IFN-γ and/or IL-2 response (2018) | Univariate | | | Multivariable | | |
|---|---|---|---|---|---|---|
| | OR | 95% CI | p value | OR | 95% CI | p value |
| Age | 0.99 | 0.96 - 1.04 | 0.89 | 1.02 | 0.97 - 1.08 | 0.35 |
| Sex | 1.67 | 0.78 - 3.63 | 0.19 | 1.57 | 0.67 - 3.75 | 0.30 |
| Baseline seropositive (HAI) | **0.23** | **0.10 - 0.51** | **0.0004** | 0.40 | 0.15 - 1.02 | 0.06 |
| Day 2 shedding | **2.87** | **1.12 - 7.82** | **0.03** | - | - | - |
| Day 7 shedding | **5.39** | **2.41 - 12.6** | **<0.0001** | **3.65** | **1.43 - 9.6** | **0.007** |

**Table S13. Impact of B/Vic shedding on B/Vic HA-specific CD4+ T-cell response (IFN-γ and/or IL-2).** Univariate and multivariable logistic regression using n = 109 children. Only participants from 2018 used as T-cell data were only generated to influenza B antigens in 2018. Excludes 17/126 participants with missing T-cell data. Day 2 and day 7 shedding variables included separately as combining into one variable (no shedding, day 2 only, day 2 and day 7) resulted in zero cell values. Only day 7 shedding taken forward to multivariable analysis. No interactions observed between variables included in the multivariable analysis. OR = odds ratio, CI = confidence interval, HAI = haemagglutination inhibition

| B/Vic HA IgA response (2018) | Univariate | | | Multivariable | | |
|---|---|---|---|---|---|---|
| | OR | 95% CI | p value | OR | 95% CI | p value |
| Age | 0.97 | 0.93 - 1.02 | 0.309 | 0.99 | 0.94 - 1.04 | 0.804 |
| Sex | 1.50 | 0.69 - 3.35 | 0.311 | 1.42 | 0.58 - 3.57 | 0.430 |
| Baseline seropositive (HAI) | **0.81** | **0.69 - 0.95** | **0.010** | 0.90 | 0.74 - 1.09 | 0.249 |
| Day 2 shedding | 1.30 | 0.55 - 3.27 | 0.553 | - | - | - |
| Day 7 shedding | **3.94** | **1.76 - 9.29** | **0.001** | 2.66 | 0.92 – 7.97 | 0.073 |
| B/Vic HA CD4+IFNg+ response | **2.42** | **1.03 - 5.71** | **0.041** | - | - | - |
| B/Vic HA CD4+IL2+ response | **2.41** | **1.05 - 5.66** | **0.039** | 1.32 | 0.48 - 3.62 | 0.679 |

**Table S14. Impact of B/Vic shedding on B/Vic HA-specific mucosal IgA response.** Univariate and multivariable logistic regression using n = 108 children. Only participants from 2018 used as T-cell data were only generated to influenza B antigens in 2018. Excludes 17/126 participants with missing T-cell data and 1/126 with poor mucosal IgA sample quality. Day 2 and day 7 shedding variables included separately as combining into one variable (no shedding, day 2 only, day 2 and day 7) resulted in zero cell values. Only day 7 shedding taken forward to multivariable analysis. B/Vic Haemagglutinin (HA) CD4+IL-2+ response taken forward to multivariable analysis due to overlap between this and the CD4+IFN-γ response. No interactions observed between variables included in the multivariable analysis. OR = odds ratio, CI = confidence interval, HAI = haemagglutination inhibition

| | | pH1N seroconversion | | | pH1 HA T-cell response | | | pH1 HA IgA response | | |
|---|---|---|---|---|---|---|---|---|---|---|
| | | Univariate | | | Univariate | | | Univariate | | |
| | | OR | 95% CI | p value | OR | 95% CI | p value | OR | 95% CI | p value |
| Age | | **0.92** | **0.85 - 0.96** | **0.03** | **0.94** | **0.89 - 0.98** | **0.01** | 0.96 | 0.89 - 1.02 | 0.207 |
| Sex | | 2.33 | 0.91 - 6.5 | 0.09 | 1.31 | 0.58 - 3.00 | 0.51 | 1.67 | 0.65 - 4.56 | 0.294 |
| Baseline seropositive (HAI)* | | - | - | - | - | - | - | - | - | - |
| Shedding | Day 2 | 3.24 | 0.58 - 24.85 | 0.19 | 1.18 | 0.41 - 3.45 | 0.76 | 1.07 | 0.24 - 4.51 | 0.929 |
| | Day 2 and Day 7 | **8.27** | **2.17 - 54.49** | **0.007** | **4.75** | **1.74 - 13.9** | **0.003** | 1.77 | 0.60 - 6.04 | 0.323 |
| pH1 HA CD4+IFNg+ response | | **4.03** | **1.47 - 12.33** | **0.009** | - | - | - | 0.61 | 0.19 - 1/74 | 0.358 |
| pH1 HA CD4+IL2+ response | | **5.31** | **1.78 - 19.7** | **0.005** | - | - | - | 1.75 | 0.60 - 5.48 | 0.314 |

**Table S15. Impact of NY15 pH1N1 shedding on pH1N1-specific immune responses.** Univariate logistic regression analyses using children immunised with 2017-18 LAIV (n = 126 in total, n = 109 in T-cell analyses, as excludes 17/126 with missing T-cell data and n = 106 in IgA analyses, as excludes 5/126 with missing pH1N1 IgA data). *Unable to use baseline haemagglutination inhibition (HAI) titre as a variable in regression due to zero cell values (e.g. No seroconverters in those that are seropositive at baseline). Multivariable analysis not undertaken as main purpose is to adjust for effect of baseline HAI titre on shedding. Shedding coded as one variable with three levels (no shedding, day 2 only shedding and day 2 and day 7 shedding). HA = haemagglutinin, IFN-γ = interferon-gamma, IL-2 = interleukin-2, OR = odds ratio, CI = confidence interval.

**<u>Percentage of participants with T-cell responses to LAIV</u>**

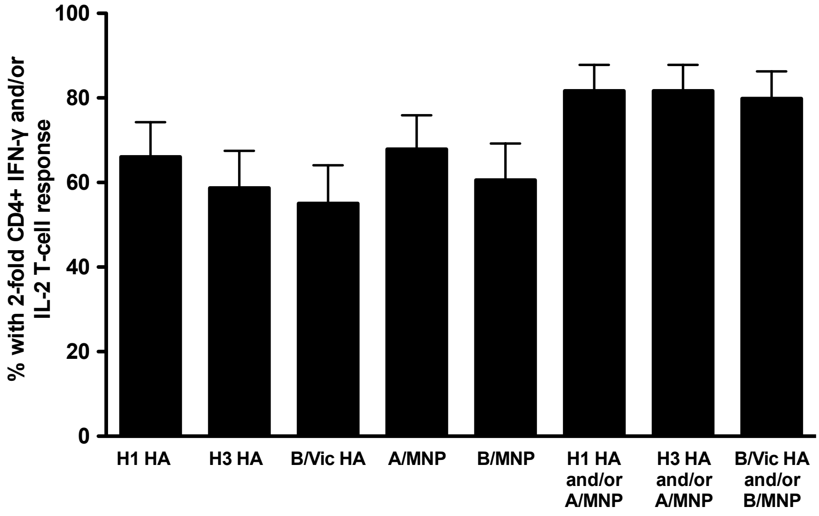

**Figure S10. Percentage of children given the 2017-18 LAIV showing a 2-fold increase from baseline to day 21 post-LAIV, in interferon-gamma (IFN-γ) and/or interleukin-2 (IL-2) producing CD4+ T-cells.** HA = haemagglutinin, MNP = matrix and nucleoprotein. Error bar represents upper 95% confidence interval.
